# Cortical propagation as a biomarker for recovery after stroke

**DOI:** 10.1101/2020.07.10.197509

**Authors:** Gloria Cecchini, Alessandro Scaglione, Anna Letizia Allegra Mascaro, Curzio Checcucci, Emilia Conti, Ihusan Adam, Duccio Fanelli, Roberto Livi, Francesco Saverio Pavone, Thomas Kreuz

## Abstract

Stroke is a debilitating condition affecting millions of people worldwide. The development of improved rehabilitation therapies rests on finding biomarkers suitable for tracking functional damage and recovery. To achieve this goal, we perform a spatiotemporal analysis of cortical activity obtained by wide-field calcium images in mice before and after stroke. We compared spontaneous recovery with three different post-stroke rehabilitation paradigms, motor training alone, pharmacological contralesional inactivation and both combined. We identify three novel indicators that are able to track how movement-evoked global activation patterns are impaired by stroke and evolve during rehabilitation: the duration, the smoothness, and the angle of individual propagation events. Results show that, compared to pre-stroke conditions, propagation of cortical activity in the acute phase right after stroke is slowed down and more irregular. When comparing rehabilitation paradigms, we find that mice treated with both motor training and pharmacological intervention, the only group associated with generalized recovery, manifest new propagation patterns, that are even faster and smoother than before the stroke. In conclusion, our new spatiotemporal propagation indicators act as biomarkers that are able to uncover neural correlates not only of motor deficits caused by stroke but also of functional recovery during rehabilitation. These insights could pave the way towards more targeted post-stroke therapies.

## 1. Introduction

Stroke is a severe disease that alters cortical processing producing long lasting motor or cognitive deficits. Treatments generally include motor rehabilitation, pharmacological therapies, brain stimulation, or combinations of them [1]. However, the functional outcome, measured as behavioral recovery, depends on multiple factors such as age, lesion size and type, edema formation or inflammation and is hardly predictable [2, 3]. One way to track recovery after stroke is by monitoring cortical activity, which is known to undergo drastic changes that have been tightly linked to structural alterations [4, 5, 6]. Previous studies have reported that stroke produces global widespread alterations in cortical activity as measured by changes in resting state functional connectivity or cortical excitability. On the one hand, electrophysiological studies have shown that stroke and recovery modulate both resting state and stimulus evoked cortical oscillations in motor areas [7, 8]. On the other hand, neuroimaging studies have shown that stroke alters the resting state functional connectivity, for example it reduces the interhemispheric correlations between motor networks, and these changes correlate with behavioral deficits [9, 10]. Furthermore, these changes in resting state functional connectivity can be used to discriminate subjects with behavioral deficits [11, 12]. This supports the idea that monitoring how cortical activity evolves over time could be used to track recovery after stroke and it could represent a powerful tool to evaluate the efficacy of stroke treatments or better yet could lead to biomarkers of functional recovery.

Here, we propose that damage and functional recovery can be tracked by monitoring the spatiotemporal properties of movement-evoked widespread activation patterns or global events. In particular, we use our recently proposed SPIKE-order analysis [13] to identify global events and to sort the participating regions from first to last (or leader to follower). Additionally, in order to characterize the spatiotemporal properties of each individual global event, we extend this method and define three propagation indicators: duration, smoothness (i.e. how ordered and consistent is the direction of the propagation), and angle.

First, we provide a characterization of global events in healthy controls. We show that these events are mostly associated with the exertion of force and that their duration and direction are modulated by different behavioral events. Then, to understand the impact of stroke on the global events we quantify the three propagation indicators in a group of acute stroke subjects. We provide evidence that acute stroke alters the propagation patterns by increasing the duration while decreasing the smoothness of the global events. To test if global events propagation patterns can be used to track recovery and to identify treatments that lead to generalized recovery, we quantify the propagation indicators in a group of subjects with untreated stroke and in three groups of subjects with different therapeutic interventions. The first therapy group received motor training alone which leads to task-specific improvement of the motor functions [14]. The second group was characterized by a transient pharmacological inactivation of the controlesional hemisphere without any improvement in the motor functions. Finally, the third therapy combined both motor training and pharmacological inactivation producing a generalized recovery of the forelimb functionality. While all treatments reduce the effect of stroke on the propagation indicators, the combined rehabilitative therapy leads to new propagation patterns defined by the shortest duration and the highest smoothness among all experimental groups.

## Methods

### Experimental set-up and data collection

In this section we provide a short overview of the data and the methods we used to analyze them. For more technical details please refer to the section “Materials and Methods” in the Supporting Information.

The aim of this study was to investigate changing propagation patterns during motor recovery from functional deficits caused by the induction of a focal stroke via a photothrombotic lesion. For this purpose, we analyzed calcium imaging signals recorded from 26 mice to compare three different rehabilitative therapies, one that used motor training alone (robot group), one based on a transient pharmacological inactivation of contralesional activity (toxin group) and one that performed both of these two treatments together (combined group).

A schematic representation of the robotic system, the M-Platform [15, 16], is shown in Fig. 1a. This system uses passively actuated contralesional forelimb extension on a slide to trigger active retraction movements that were subsequently rewarded (up to 15 cycles per recording session). The effect of the motor activity was monitored via the discrete status of the slide and by recording the force the mice applied to the slide. As a measure of the neural activity itself we performed wide-field calcium imaging over the affected hemisphere, from the somatosensory to the visual areas. Selecting a region of interest of 2.16 × 3.78 mm and spatially downsampling by a factor 3 resulted in calcium images of 12 × 21 pixels of size 180 *μ*m. These are the signals that we analyzed.

**Figure 1:**
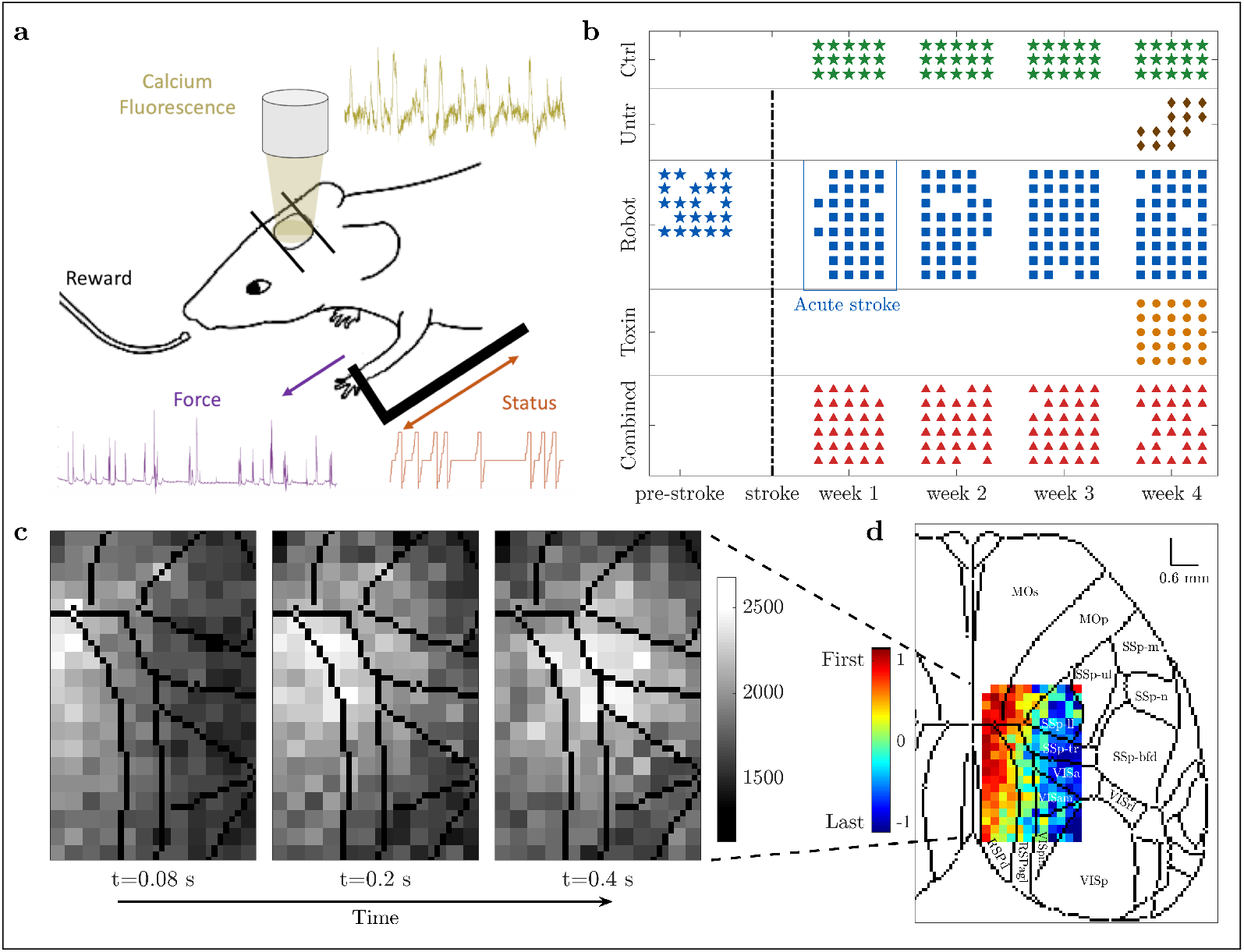
Experimental set-up and data collection. **(a)** Motor-training of mice was performed on the M-Platform, which uses a movable robotic slide for a retraction movement of the left forelimb. Motor activity was monitored via the discrete status of the slide (orange) and the force applied by the mouse to the slide (purple). Meanwhile, the cortical activity (yellow) was recorded using wide-field calcium imaging. **(b)** The control (green) and combined treatment (red) group performed four weeks of daily training; for the robot (blue) group we additionally recorded one week before stroke induction (typically five sessions per week). Untreated (brown) and toxin (orange) groups only performed one week of daily training starting four weeks after the lesion. Star symbols refer to the healthy condition. **(c)** Calcium imaging sequence of cortical activation, superimposed on contours of brain regions according to the standard atlas. **(d)** Propagation pattern, from leader (red) to follower (blue), of the event depicted in (c). Color coding is based on the SPIKE-order (for details see section “SPIKE-order” in “Materials and Methods” (Supporting Information)). For a complete list of brain regions and their acronyms see Fig. S1 in the Supporting Information.

The 26 mice were divided into five groups: control (3 mice), untreated (4 mice), robot (8 mice), toxin (5 mice), and combined treatment (6 mice). The healthy controls had no stroke induced but underwent four weeks of motor training on the M-Platform. The untreated mice performed one week of motor training starting 25 days after injury. The toxin group, which received Botulinum Neurotoxin E (BoNT/E) injection on the contralesional hemisphere right after the photothrombotic damage, also performed one week of motor training starting 25 days after injury. Both robot and combined treatment mice underwent physical rehabilitation on the M-platform for four weeks starting five days after injury. For toxin and combined treatment mice the pharmacological inactivation of the primary motor cortex in the contralesional hemisphere was carried out in order to counterbalance the hyperexcitability of the healthy hemisphere.

The recording schedule for all five groups is shown in Fig. 1b. Apart from the data acquisition during the shared training regime, 5 out of 8 robot mice were also recorded for one week before the stroke (pre-stroke condition). We have confirmed that this pre-stroke condition shows no statistical difference, both qualitatively and quantitatively, with the first week of recordings of the control group, as could be expected for two groups of healthy mice (for details see Fig. S2 in Supporting Information).

Fig. 1c displays a sequence of snapshots of the calcium activity over time. The three images illustrate one pull of the slide by the contralateral forelimb of a control mouse, from the activation of the average calcium activity (left) via its maximum (middle) to its tail end (right). The Supporting Information contains a movie that covers all 36 frames for this one training cycle.

The central method of this study was a mapping of this sequence of snapshots into the propagation pattern shown in Fig. 1d. In this matrix the order of activation of the individual pixels was color-coded from red (earliest) to blue (latest). We superimposed this order matrix on the standard atlas of brain regions [17] which illustrates that the recording area covers the primary motor area M1 (the location of the lesion), the primary somatosensory area, and the primary visual cortex, as well as the retrosplenial cortex.

### Event identification, propagation analysis, and definition of three propagation indicators: duration, angle, and smoothness

Here we explain our use of the SPIKE-order framework [13] to identify global events and assess the propagation of activation within these events (compare Fig. 1d). In particular, we focus again on one individual global event to illustrate how we characterize the detailed activation patterns with three propagation indicators: **duration**, **angle**, and **smoothness**.

In Fig. 2 we show the activity during a complete recording session of one mouse. Figure 2a depicts the status, a discrete codification of the current phase of the passive extension and active retraction cycle, e.g. position of the slide and acoustic go and reward cues. Most relevant here are the marked times of the pull completions which typically correspond to peaks in the force applied to the slide (Fig. 2b, force events are marked at threshold crossings) but already a quick look at the event numbers shows that the mapping is not perfect. Indeed there are typically more force events than rewarded pull completions (20 force versus 13 status events in this case). The same peaks, and some more, are also present in the calcium signal computed by averaging the fluorescence signal over all pixels (Fig. 2c, here 23 calcium events are marked at threshold crossings).

In the next step we looked at all individual pixels but here we do not show all the traces but just small time markers denoting the time of their threshold crossings (Fig. 2d). This is very similar to a rasterplot showing in each row the spike train of one individual neuron and accordingly we here follow this terminology and call the threshold crossings of individual pixels ‘spikes’. The first thing to notice is that while there are a few spikes in the background (in black), by far most of the spikes are part of global events (in color) matching the peaks in the average calcium activity shown right above. To automatically identify these global events and sort the spikes within these events from leader to follower we used the SPIKE-order framework proposed in [13].

**Figure 2:**
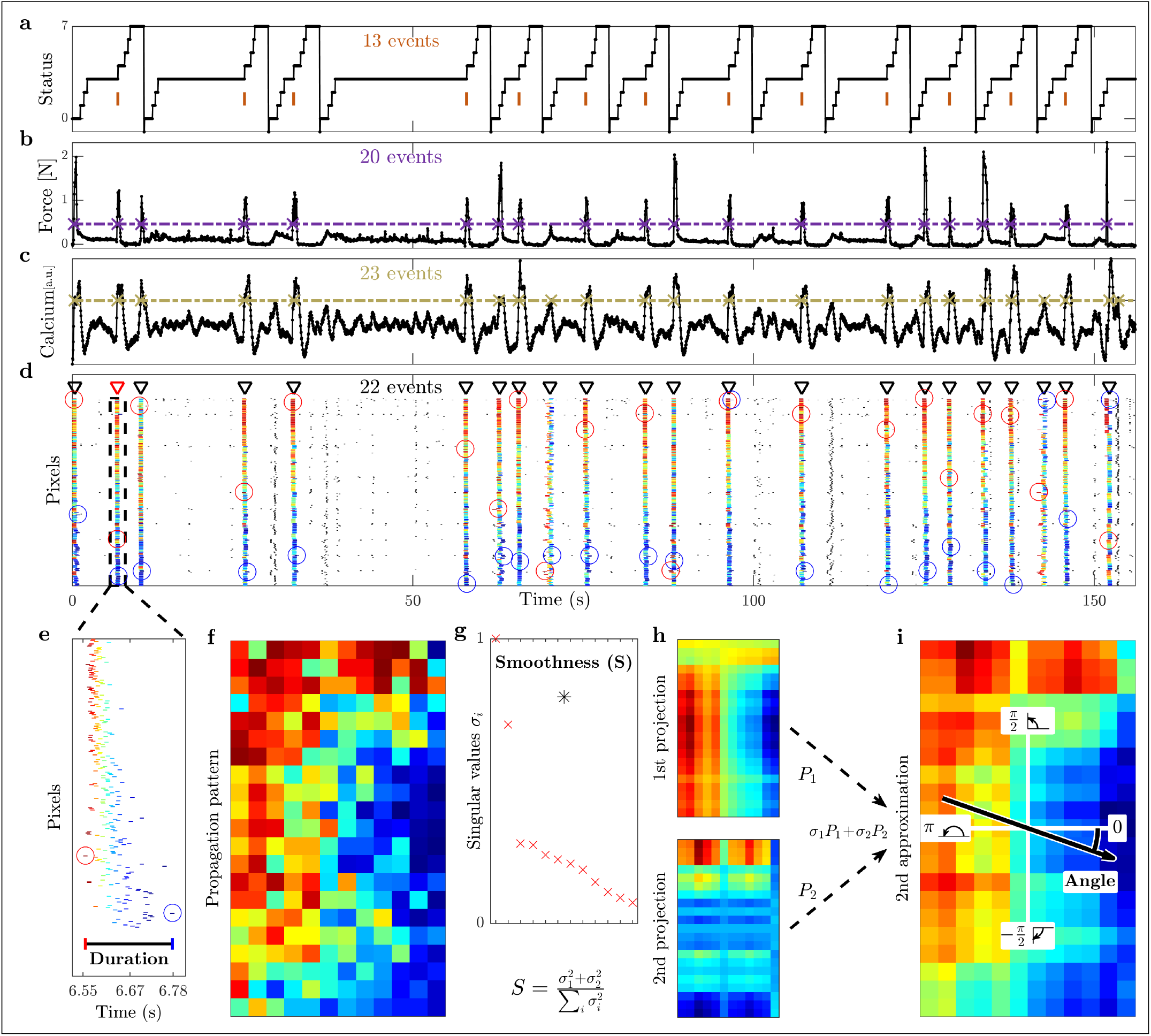
Event identification, propagation analysis, and definition of three propagation indicators: duration, smoothness, and angle. **(a)** Status of the robotic slide. The longer horizontal plateaus at status 3 correspond to the time interval during which the mouse is allowed to retract the slide. The orange bars refer to the time when the pull is completed and the mouse receives its reward. **(b)** Force applied by the mouse during the retraction movement. **(c)** Average calcium signal over all pixels. The purple and yellow dashed lines below refer to the threshold used to identify the force and global calcium events, respectively. **(d)** Raster plot obtained from the threshold crossings of individual pixels versus time. Triangle indicators show the global events identified during this session, and the red triangle with dashed box marks the event analyzed in the remaining panels below. The first (last) spike of each global event is marked by a red (blue) circle. Within each subplot (a)-(d) we state the number of the respective events / threshold crossings. **(e)** Zoom of this selected event. The **duration** is defined as the interval from the first to the last spike within the event. **(f)** The propagation matrix is obtained by projecting the relative order of these threshold crossings onto the 2D-recording plane. **(g)** Singular values (red) vs. order of approximation, obtained by means of singular value decomposition (SVD). The **smoothness**(black asterisk) measures the quality of the second order approximation. **(h)** First and second approximations of the propagation matrix and **(i)** second order approximation as their weighted sum (cumulative). The **angle** of the propagation, defined relative to the horizontal axis.

After some initial denoising we first used an adaptive coincidence detector [18] to pair spikes such that each spike is matched with at most one spike in each of the other pixels. By means of the symmetric and multivariate measure SPIKE-Synchronization [19] we filtered out all spikes which were not coincident with spikes in at least three quarters of the other spike trains. To the global events that remained we applied the asymmetric SPIKE-order indicator [13] which quantifies the leader-follower relationship between pairs of spikes. For each event the SPIKE-order is then color-coded from leader (red) to follower (blue). Finally, we used the scalar Synfire Indicator [13] to also sort the spike trains in the rasterplot from overall leader to overall follower. Since the sorting takes into account all global events, the first spike trains contain more leading spikes (red) and the last spike trains more trailing spikes (blue).

In Fig. 2e we zoom in on the fourth global event of the rasterplot. Here we define the first propagation indicator, the event **duration**, as the time from the first to the last spike of this event. The propagation matrix of this specific event, obtained by projecting the color-coded relative order of the spikes onto the pixels of the 2D-recording plane, is shown in Fig. 2f. We then used singular value decomposition (SVD) to calculate the two remaining propagation indicators.

SVD [20] searches for spatial patterns by decomposing the propagation matrix into three simple transformations: a rotation, a scaling along the rotated coordinate axes and a second rotation. The scaling is a diagonal matrix which contains along its diagonal the singular values of the propagation matrix. In Fig. 2g we depict the (sorted) singular values *σ_i_* (rescaled to the highest value). We also show the value of the second propagation indicator, the **smoothness***S*, which is defined as the relative weight of the first two projections, their sum divided by the sum of all singular values. Backprojecting the (sorted) singular values one at a time resulted in various orders of approximations for the original propagation matrix. The first two such projections are displayed in Fig. 2h and the second order approximation, their weighted sum, is shown in Fig. 2i. From the weighted average of the mean gradients of these first two projections we calculated the third propagation indicator, the **angle** of the main propagation direction. Note that the smoothness quantifies how well the approximation using only the first two singular values captures the full spatiotemporal pattern obtained by considering all singular values. This can be verified visually by comparing the second approximation (Fig. 2i) with the original propagation matrix (Fig. 2f).

A comparison of Fig. 2d and Fig. 2c clearly shows that all global events in the rasterplot can easily be matched with a peak in the average calcium trace, in fact, these are basically two equivalent ways to visualize a peak of global calcium activity. However, while the vast majority of global events are in close proximity of a peak in the force, not all of them are. This we can use to categorize the global events into two groups: Force (**F**) and non-Force (**nF**). Among the Force events we can distinguish two kinds of events, a few of them occur during the passive extension of the arm by the slide (Passive, **Pass**) but most of them do not, i.e., they occur during the active retraction phase (Active, **Act**). Finally, among those active events we can differentiate between movements which lead to a completion of the forelimb retraction and therefore are rewarded (Reward Pulling, **RP**) and movements which are not completed and thus not rewarded (non-Reward Pulling, **nRP**). The Reward Pulling events are the ones that correspond to the vertical markers in the status trace of Fig. 2a.

The overall categorization can be visualized by means of this branching structure:

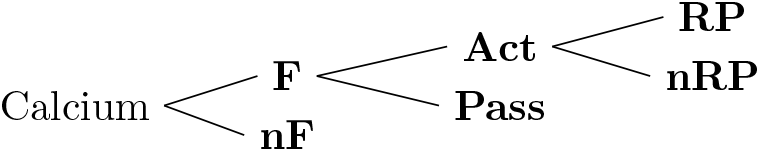

## Results

In this study, we sought a biomarker for functional recovery after stroke in the neural activity propagation. To this aim, we used global event propagation analysis to investigate the spatiotemporal features of neuronal activation over the dorsal cortex. We compared the propagation patterns of healthy mice versus acute stroke and of untreated versus treated mice. Here we looked at three different treatments (robot, toxin, and combined, see Fig. 1b). We decided to investigate three main indicators (duration, smoothness, and angle of the propagation) to account for both temporal and spatial propagation characteristics (compare Fig. 2). To identify common brain dynamics associated with similar behaviors, we dealt with each type of event separately.

### Cortical propagation features discriminate event types in healthy mice

We first wondered if neuronal activation in awake healthy mice involved a large region of the cortex and if these global events were related to specific classes of behavioral events in our experimental paradigm. The results in Fig. 3 focus on two of the three indicators introduced in Fig. 2 and they refer to one healthy mouse during all sessions (four weeks, five days per week, see Fig. 1b). Three different example patterns with increasing smoothness and varying angle are depicted in panels a-c. For low smoothness values, the identified propagation pattern looks random, thus not displaying a clear directionality (panel a), and therefore measures for angle and duration are less meaningful in those cases. On the other hand, high smoothness corresponds to clear patterns (panels b-c show two cases of high smoothness but orthogonal directionality). Panels d-f show scatter plots of smoothness against angle for different event types, together with the marginal histograms. While narrowing down the type of event does not reduce the whole range of values, the marginal distributions of the angle and smoothness converge to a peak distribution for the angle centered in 0.46, and to a distribution with mean 0.68 for the smoothness.

**Figure 3:**
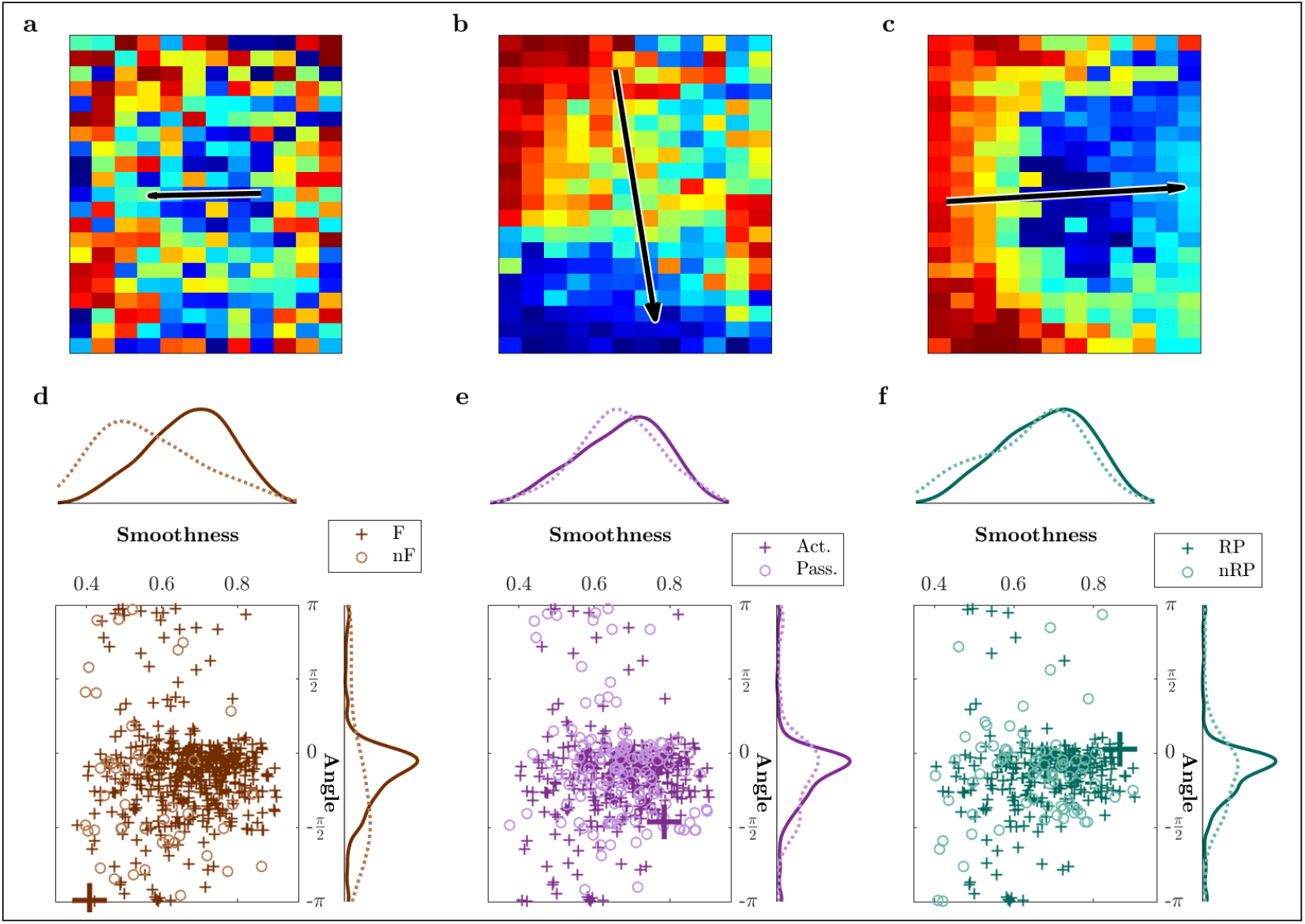
Propagation pattern varies across event types in healthy mice. **(a-c)** Three different patterns with increasing smoothness and varying angle. Note that the length of the arrow is proportional to the smoothness. **(a)** Low smoothness identifies a poor propagation pattern without any proper directionality. **(b)** High smoothness identifies a clear pattern and corresponding directionality. **(c)** High smoothness and horizontal propagation in contrast with (a) low smoothness and opposite horizontal direction, and different from (b) which has still high smoothness, but an almost orthogonal angle of *π/*2. **(d-f)** Scatter plots and histograms of smoothness and angle of propagation for all events. The events whose propagation patterns are shown in (a)-(c) are highlighted by larger markers. — Brown colors refer to force (F) and non-force (nF) events, active (Act) and passive (Pass) events are in purple, reward pulling (RP) and non-reward pulling (nRP) events in green. All plots include the events from all the sessions for one healthy mouse.

Next, we performed a detailed quantitative analysis of all three indicators (duration, smoothness and angle) for all three healthy mice along four weeks of motor training on the M-Platform (control group, Fig. 4). The dependency of the three indicators was evaluated both over time and with respect to different event types. In the beginning we analyzed if the occurrence of the global events was associated with the application of forces or if it was unrelated. For the four weeks of recordings in healthy mice the variation of the number of events divided by type is depicted in Fig. 4a. Most of these events occurred when the mouse applied a force to the handle (1568, corresponding to 87% of the total number of events). Furthermore, most of the force events (1147) occurred when the mouse was actively pulling the handle (73% of force events, 64% of the total), and 779 of those corresponded to reward pulling events (68% of active events, 43% of the total).

**Figure 4:**
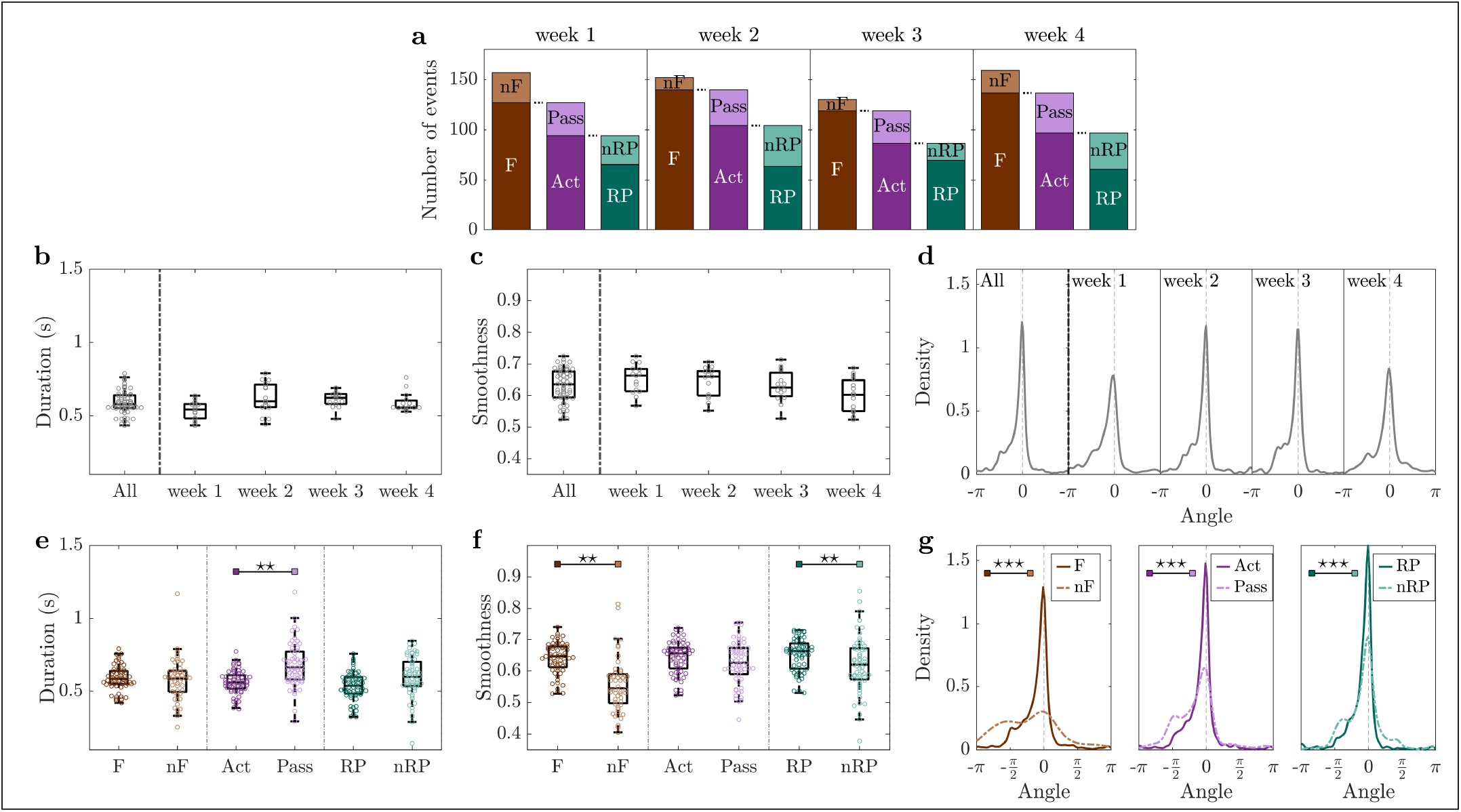
Cortical propagation features discriminate event types in healthy mice. **(a)** Mean number of events per mouse per week, partitioned by type of event. **(b-d)** Duration, smoothness, and angles distribution are preserved over weeks. **(e)** Event duration is a marker for discriminating active (Act) and passive (Pass) events. **(f)** Smoothness discriminates force (F) and non-force (nF) events, as well as reward pulling (RP) and non-reward pulling (nRP) events. **(g)** Narrowing down the type of event leads to more directed propagation patterns. — Duration and angle are weighted by smoothness. Markers in (b,c,e,f) refer to the average value per day. Within each box in (b-f), the central mark indicates the median, and the bottom and top edges of the box indicate the 25^*th*^ and 75^*th*^ percentiles, respectively. Control group n=3 mice. P-values of statistical tests in Table S2 (Supporting Information), “*\**” refers to difference in variance.

Global events last 0.59 ± 0.04 s with a smoothness of 0.63 ± 0.03, and they propagate along angles −0.3 ± 0.4, see subplot “All” of Fig. 4b-d. The little variation in duration (Fig. 4b), smoothness (Fig. 4c) and angle (Fig. 4d) implies high coherence of the parameters of spatiotemporal propagation over weeks. This suggests that longitudinal motor training in healthy mice does not alter the number of events or the propagation patterns.

Results for duration, smoothness and angle were then analyzed looking at specific event types. Fig. 4e shows the event duration for the three consecutive subdivisions of the event types. Further specifying the type of event leads to shorter and shorter propagations with the shortest average duration being obtained for reward pulling events. The same argument can be made for the smoothness (Fig. 4f); reward pulling events display the highest smoothness, on average, among all the other events. Fig. 4g shows the distributions of the angle; the propagation becomes more directed when narrowing down the type of event. The difference between event types in the angle distribution reduces as well. Interestingly, also the bimodal nature of the distribution is attenuated. Specifically, the peak at −*π/*2 (see first plot in Fig. 4g) is initially caused by non-force events, then within force events it stands out in the passive cases, finally it is predominant in the non-reward pulling events. This suggests that task-specific events, such as reward pulling, are characterized by consistent propagation patterns.

The patterns observed in healthy mice are characterized by different spatiotemporal features when comparing force and non-force events in terms of smoothness (p=0.001) and angle of propagation (p=10^*−*10^), and when comparing active and passive events in terms of duration (p=0.002) and angle (p=10^*−*9^). Also between rewarded and non-rewarded pulls events the observed patterns display different characteristics when comparing smoothness (p=0.001) and angle (p=10^*−*8^), see Fig. 4e-g and Table S2 (Supporting Information) for the p-values.

In summary, in healthy mice there is a high coherence of the parameters of spatiotemporal propagation over weeks, suggesting that a simple motor task alone does not change the duration, smoothness, and angle of events. Moreover, the investigated spatiotemporal propagation indicators discriminate between different event types when a specific characteristic is taken into account, i.e., force versus non-force, active versus passive, reward versus non-reward pulling.

### Acute phase after stroke is characterized by an increase of event duration

We pondered how the spatiotemporal propagation indicators were altered by cortical injury, and thus looked at the cortical activation events as associated to classes of behavioral events in the first week after stroke (called acute stroke). Moreover, we compared these results with the first week of recordings on healthy mice (Fig. 5) which consists of the first week of recordings of the control group and the pre-stroke week in the robot group (see Fig. S2 in the Supporting Information).

**Figure 5:**
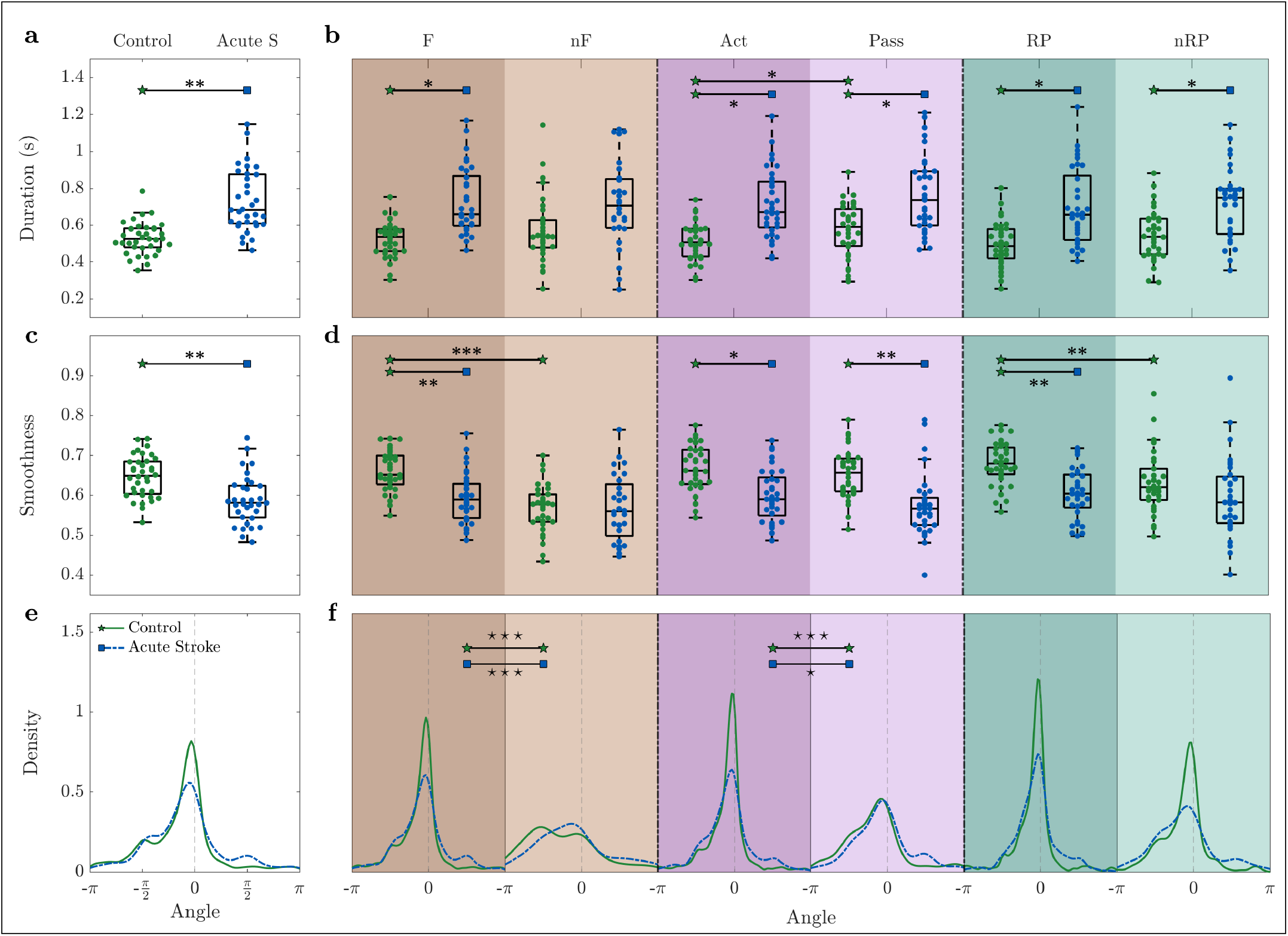
The acute phase (one week after stroke) is characterized by an increase of the event duration and decrease of smoothness. During the acute phase, **(a-b)** the event duration is increased, **(c-d)** smoothness is decreased, and **(e-f)** the direction of propagation is more spread. — Duration and angle are weighted by smoothness. Markers in (a-d) refer to the average value per day. Within each box in (a-d), the central mark indicates the median, and the bottom and top edges of the box indicate the 25^*th*^ and 75^*th*^ percentiles, respectively. Control n=8 mice, Acute Stroke n=8 mice. See Table S3 (Supporting Information) for p-values of statistical tests, “*\**” refers to difference in variance and “*” refers to difference in mean.

When looking at all events together, differences can be appreciated for the duration (p=0.003) and the smoothness (p=0.007) even without further splitting the events into specific types. In particular, the duration increases (Fig. 5a) and the smoothness decreases (Fig. 5c) during the acute phase. The angles distribution for the acute stroke group exhibits a flatter distribution and two secondary peaks in −*π/*2 and *π/*2, indicating the presence of a more heterogeneous pool of events (Fig. 5e).

When splitting the results into event types (Fig. 5b,d,f), a common tendency for duration, smoothness and angle is that the stroke condition attenuates differences of the indicators between event types. For both duration and smoothness (Fig. 5b,d), while the control group presents significant variations with respect to the type of event (Act-Pass p=0.04 for duration, and F-nF p=10^*−*10^ and RP-nRP p=0.008 for smoothness), the acute stroke group is characterized by smaller fluctuations in the mean value (Act-Pass p=0.059 for duration, and F-nF p=0.082 and RP-nRP p=0.51 for smoothness). For the duration (Fig. 5b), significant differences in mean between healthy and stroke mice are manifested for force (F, p=0.026), active (Act, p=0.034), passive (Pass, p=0.035), reward pulling (RP, p=0.014), and not reward pulling (nRP, p=0.028) events. Regarding the smoothness (Fig. 5d), significant differences can be found in the same event types as for duration, non rewarded pulls (nRPs) excluded (p=0.004, 0.01, 0.002, 0.005). For the angle (Fig. 5f), the dissimilarities in the control group between event types are preserved in the acute stroke condition (F-nF and Act-Pass, control p¡0.001 and robot p¡0.05).

Altogether, our three spatiotemporal propagation indicators are able to distinguish between healthy and stroke mice. The acute phase after stroke leads to more heterogeneous events characterized by longer duration, lower smoothness, and flattened distributions of the angle. Moreover, in contrast to the healthy condition, for the acute stroke condition cortical propagation features are not able to discriminate event types.

### Combined rehabilitative treatment induces short duration and high smoothness of cortical propagation patterns

Our previous results comparing motor, pharmacological and combined therapy after focal stroke demonstrated that only a rehabilitation protocol coupling motor training with reversible inactivation of the contralesional cortex was able to promote recovery in forelimb functionality [14]. Here we first analyzed the propagation patterns in spontaneously recovered mice (untreated stroke group) one month after the lesion (Fig. S3 in the Supporting Information). Results show that untreated animals present only minor modulations in the propagation features with respect to the acute phase. The only statistical difference between the two groups is found for smoothness, which is lower for the untreated group, maintaining the same trend induced by the stroke in the acute phase (compare Fig. 5).

We hypothesized that rehabilitative treatments alter the spatiotemporal propagation patterns, and especially reverse the trend observed in the acute stroke phase. In particular, given the fact that the combined treatment is the only treatment that leads to a generalized recovery (see Fig. S4 in the Supporting Information), we wondered if we can find for this treatment spatiotemporal features of cortical propagation that reflect this unique success. We thus compared the spatiotemporal propagation indicators in treated animals with spontaneously recovering mice (untreated stroke group). In detail, we evaluated the consequences on the propagation patterns in mice treated with motor training alone (robot group), pharmacological silencing of the homotopic cortex alone (toxin group) or a combination of both (combined treatment group), see Fig. 6.

**Figure 6:**
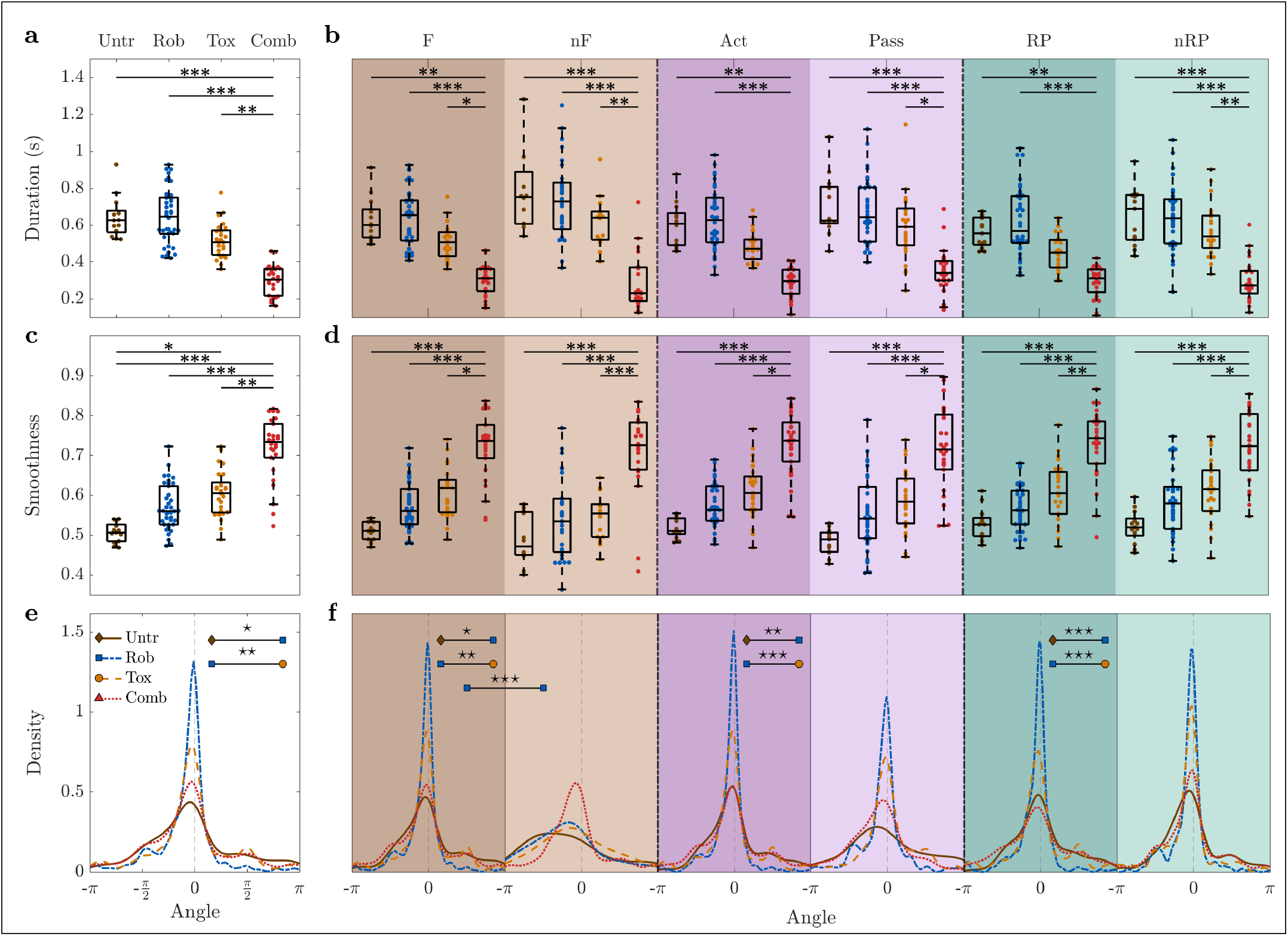
Combined treatment group is characterized by higher smoothness and shorter duration than any other group. For all types of event, combined treatment group events are the shortest **(a-b)** and their smoothness is the highest **(c-d)**. For the combined treatment group the distribution of the angles does not vary depending on the type of event **(e-f)**. — Duration and angle are weighted by smoothness. Markers in (a-d) refer to the average value per day. Within each box in (a-d), the central mark indicates the median, and the bottom and top edges of the box indicate the 25^*th*^ and 75^*th*^ percentiles, respectively. Untreated stroke group n=4 mice, robot group n=8 mice, toxin group n=5 mice, combined treatment group n=6 mice. For P-values of statistical tests see Table S4 (Supporting Information), “*\**” refers to difference in variance and “*” refers to difference in mean.

Indeed, the most striking result emerging from this analysis is that the combined treatment group is greatly separated from all the other groups. When looking at all events together (Fig. 6a), the events of the combined treatment group display a shorter duration compared to untreated (p=10^*−*4^), robot (p=10^*−*5^), and toxin (p=0.006). The biggest differences can be observed for combined treatment versus untreated and robot groups; they are significant not only for all events together but also for each event type separately (all p<0.006 and all p< 10^*−*4^, respectively), see Fig. 6b.

Among all the characteristics investigated, the marker that distinguishes most clearly between the combined treatment group and all the others is smoothness. The events of the combined treatment group display a greater smoothness compared to the untreated (p=10^*−*5^), robot (p=10^*−*4^), toxin (p=0.003) groups (Fig. 6c). Again, this statement applies not only to all events together, but also to each event type individually (all p< 10^*−*4^ for untreated and robot, all p<0.023 for toxin), see Fig. 6d.

Dissimilarities in the angle distributions are able to capture the difference between the robot group and both untreated and toxin groups (Fig. 6e,f). Note that all three indicators of the combined treatment group appear to be consistent when distinguishing different event types, meaning force, non-force, active, passive, reward and non-reward pulling events display similar average values. The complete list of p-values can be found in Table S4 (Supporting Information).

Additionally, we evaluated if propagation features were restored to pre-stroke levels or if the treated condition ended in a new state (Fig. S5-6 in the Supporting Information). We analyzed longitudinal data starting from the second week of recording up to one month after stroke, by comparing motor treated (robot and combined) with healthy mice (control group), see Fig. S5. Results show that propagation features averaged over the training weeks were partially restored to pre-stroke levels for the robot group (no statistical differences were found with the control group), while the combined treatment group ended in a new state with very different values from both pre-stroke and motor-treated conditions. In details, the events of the combined treatment group display shorter duration (p=10^*−*4^, p=10^*−*6^) and a greater smoothness (p=0.025, p=10^*−*4^) compared to the control and robot groups. For a complete list of p-values refer to Table S8 (Supporting Information). When looking at the temporal evolution of the recovery (Fig. S6), in the weeks that follow the stroke the robot group shows a significant variation: the smoothness is lower and the duration is longer. This variation decreases over time in both cases. While the duration reaches again values comparable to healthy mice already after the second week of training, the difference in smoothness seems to oscillate without stabilizing. Interestingly, the rehab group presents a behavior qualitatively comparable to the control group, i.e., a stable trend, but with very different values.

In summary, the combined treatment group is significantly different from all the other groups. In particular, it is characterized by the shortest duration and the highest smoothness. These differences can not only be observed for all events together, but also for all event types (force, non-force, active, passive, reward pulling, non-reward pulling) separately. Once more, this confirms that the combined therapy, the only one leading to generalized recovery, is associated with a state different from pre-stroke conditions which shows new and unique propagation features.

## Discussion and conclusions

In this study we employed an improved version of our recently proposed SPIKE-order analysis [13] to sequences of wide-field fluorescence calcium images from the dorsal cortex of awake behaving mice. We defined three propagation indicators that characterize the duration, the angle of propagation and the smoothness of movement-evoked global events. This new way of quantifying variations in the spatiotemporal propagation patterns allowed us to track damage and functional recovery following stroke.

We found that in healthy mice all three indicators of spatiotemporal propagation display a very high degree of consistency over time. For animals with acute stroke the propagation patterns of the global events through the injured hemisphere are altered. The most prominent consequence is a large increase in global event duration and a decrease in smoothness over the ipsilesional hemisphere. We compared spontaneous recovery with three different rehabilitation therapies, motor training, transient pharmacological silencing of the homotopic cortex and a combination of both. While all of these treatments have an impact on the spatiotemporal propagation patterns, the combined therapy, promoting a generalized recovery of forelimb functionality, leads to an increased propagation efficacy, different from pre-stroke conditions, with very fast and smooth patterns.

### Comparison with existing methods

Most analysis tools for wide-field optical images perform simple correlation and time lag analysis (Pearson correlation and phase synchrony), which are window- and not event-based as in this study (e.g. [21, 22, 23]). Directionality is explored by using Granger causality in [24], which is dependent on a priori selection of regions of interest (ROIs). Since commonly used analyses tools are based on averaged activity under resting state, important information on single events is missing. This is especially important when considering motor evoked activity, which is directly associated with the execution of single movements (active forelimb pulling in our case). The detailed propagation analysis we showed here would not be possible with the widely used optical flow techniques [25, 26] which instead focus on velocity vector fields and their complex patterns (e.g. sources and sinks) but do not deal explicitly with temporal order. Our methods used here contain some similarities but also crucial differences with a very recent analysis performed on cortical slow waves in anesthetized mice [27]. On the one hand, both algorithms apply exactly the same criteria of unicity and globality in the identification of the spatiotemporal patterns. On the other hand, the core of our global event detection is automated spike matching (via adaptive coincidence detection) while the wavehunt pipeline relies on an iterative procedure to cut the time series into distinctive waves. Apart from the different types of data, the two studies are also complementary in scope: while the focus of [27] lies on the excitability of the neuronal population, the dominant origin points and the velocity of the slow waves, we perform a thorough investigation of the spatial propagation pattern of each global event.

As a final remark on the methods, we would like to stress that the approach used here is universal and could easily be adapted to functional techniques (including electroencephalography and functional Magnetic Resonance Imaging) in many other clinical settings. For example, it could be extended to disorders of the central nervous system similarly associated with alterations in the spatiotemporal propagation of brain activity, from traumatic brain injury to autism.

### Cortical propagation features in healthy mice

We first tested the discrimination capabilities of this approach for different classes of behavioral events and used it to characterize global events over most of the dorsal cortex. Results show that under our experimental paradigm, in all conditions, global events are occurring predominantly when the mouse is actively applying force during either active retraction or passive extension of the affected fore-limb. Angle, duration and smoothness of the global events change with behavioral event type (e.g. if the event is associated with the application of force or not) in healthy subjects. This finding is in line with a recent study showing different propagation patterns across the cortex for mice engaged in a visual task depending on the type of the behavioral event (active vs passive, hit vs misses, ipsilesional vs controlesional) [28].

The drastic change observed in the angle distribution of global events between force and non force events implies that activity propagates from medial to lateral regions. This is in accordance with previous findings based on space-frequency single value decomposition analysis showing that at the naive stage, the activity propagated from retrosplenial cortex in a radial direction [29]. The mediolateral propagation of the global events suggests the progressive involvement of the retrosplenial cortex during the exertion of the reward pull. Indeed, it has been previously reported that retrosplenial cortex is more correlated with sensory cortices during locomotion vs quiescence [30] suggesting the presence of a network switch to allow the processing of sensory information during locomotion. In this view, the higher accumulation of angles at 0 degrees observed when comparing force vs non force events could represent the hallmark of such network switch. Interestingly, a similar propagation pattern has been observed applying optical flow analysis to calcium imaging over the same cortical areas when windowing cortical activity around hippocampal sharp wave ripples during sleep [31] and corresponds to one of the two major propagation patterns observed during slow wave sleep [32]. Our findings extend these results to the awake condition during motor execution.

### Acute phase after stroke

Stroke strongly affects the spatial propagation within the cortex during the execution of a pulling task. The analysis of spatiotemporal propagation patterns evoked by stimulation or voluntary movements represents a fundamental means to investigate functional remapping in order to better understand post stroke reorganization. In our study, acute stroke is characterized by less coherent direction of the propagation, lower smoothness, and longer duration than healthy animals. Also, cortical propagation properties are very heterogeneous across global events and the differences between behavioral event types are lost. In a previous work Murphy and collaborators [33], by applying intrinsic optical signal and fluorescence imaging, described modifications in spatiotemporal propagation elicited by sensory forelimb stimulation both in acute and chronic phase after stroke in the forelimb cortex. In agreement with Brown et al., who observed during the acute phase an increase of time to peak cortical signal evoked by sensory stimulation, our results revealed an increment of duration of motor-evoked cortical response, showing a delayed activation of cortical regions neighboring the stroke core due to the damage. The comparison between these results reveals that though applying opposite approaches (i.e. bottom-up for sensory stimulation and top-down for motor task execution) a similar cortical response was observed. Moreover, an fMRI study in the acute phase by Dijkhuizen and colleagues [34] showed that the stimulation of the unpaired forelimb induces a small response detected in the ipsilesional hemisphere in M1 and sFL cortex and in more distal regions both in rostral and caudal direction. A similar observation was made by Harrison and collaborators [35] revealing that motor maps were more diffuse after motor-targeted stroke during sensory stimulation, with a decrease in correlation between neighboring pixels. The diffuse activation in response to forelimb stimulation observed in those previous works is in agreement with our results that reveal the absence of a clear pattern of cortical propagation, as highlighted by low smoothness, in the acute phase after stroke.

### Comparison of different rehabilitation paradigms

Different rehabilitative paradigms result in different rehabilitative outcomes; in particular, not all treatments promote generalized recovery of forelimb functionality. In this study, we purposely selected three rehabilitative treatments (robot, toxin, and combined treatment) that all induced changes in the neural signals but only the combined treatment leads to generalized recovery. Combined rehabilitation profoundly altered the propagation of global events as compared to both healthy (control) and post-stroke single treated (toxin and robot) mice. The differences in the propagation features induced by stroke in the acute phase decline during weeks of motor training in the robot group, in fact already by the second week of training duration reaches values comparable to healthy mice. Also the smoothness was on average comparable to healthy mice after robotic training. Our findings on the chronic phase of the robot group are in agreement with what we observed in [36] where repetitive motor training induced a task-dependent spatial segregation similar to healthy mice though unaccompanied by functional recovery [14]. Each treatment had an impact on separate propagation features. In fact, while during the chronic phase only robot mice showed significant differences in the direction of cortical propagation (angle) compared to untreated stroke mice, the toxin group had peculiar (and significantly different) features in smoothness. Combination of the two treatments results in profound global changes in all indicators, with new features completely different from all the other groups. More in detail, combined treatment mice show a decrease of duration and greater smoothness with respect to control and robot mice, indicating the emergence of faster and more directed patterns of propagation. Such temporally compressed and reliable cortical activity sequences may be associated with a more effective trigger of subcortical movement machinery [29].

In addition, the substantial increase in smoothness after combined rehabilitation finds a nice correlate in the segregation of motor representation illustrated in preclinical [6] and clinical studies [37]. In these works, improved motor functionality induced by post-stroke combined rehabilitation is associated with a more focused brain activation during the execution of a motor task [36]. Importantly, generalized recovery of forelimb functionality in combined treatment mice (see [14, 6]) is not necessarily associated with recovery of pre-stroke spatiotemporal propagation features. Indeed, the results on all motor-evoked spatiotemporal propagation indicators suggest that the combination of contralesional inactivation and motor training acts towards the establishment of new propagation patterns rather than the restoration of pre-stroke features.

In summary, our detailed spatiotemporal analysis of global activation patterns during longitudinal motor training provides a powerful non-invasive tool to quantify the success of different state-of-the-art rehabilitation paradigms. The propagation-based biomarkers deliver new and unforeseen information about the brain mechanisms underlying motor recovery which could pave the way towards a much more targeted post-stroke therapy.

## Source Codes

The main method used here, SPIKE-Order, is implemented in three free and publicly available software packages. Results in this study were obtained using cSPIKE^2^ (Matlab command line with MEX-files). A Matlab-based graphical user interface, SPIKY^3^ [19], and a Python library, PySpike^4^ [38], are available as well.

## Acknowledgments

We thank Shih-Chieh Lin and Cristina Spalletti for their reading of the manuscript and their insightful comments. This project has received funding from the H2020 EXCELLENT SCIENCE - European Research Council (ERC) under grant agreement ID n. 692943 BrainBIT and from the European Union’s Horizon 2020 Research and Innovation Programme under Grant Agreement No. 785907 (HBP SGA2) and No. 945539 (HBP SGA3). The authors declare that they have no known competing financial interests or personal relationships that could have appeared to influence the work reported in this paper.

## Footnotes

2 http://www.fi.isc.cnr.it/users/thomas.kreuz/Source-Code/cSPIKE.html

3 http://www.fi.isc.cnr.it/users/thomas.kreuz/Source-Code/SPIKY.html

4 http://www.pyspike.de

