## Supplementary Information for "Cortical propagation as a biomarker for recovery after stroke"

### Materials and Methods

*Experimental design*

*Mice*

All experimental procedures were performed in accordance with directive 2010/63/EU on the protection of animals used for scientific purposes and approved by the Italian Minister of Health, authorization n.183/2016-PR. Mice were housed in clear plastic cages under a 12 h light/dark cycle and were given ad libitum access to water and food. We used a transgenic mouse line (C57BL/6J-Tg(Thy1GCaMP6f)GP5.17Dkim/J, referred to as GCaMP6f mice) expressing a genetically-encoded fluorescent calcium indicator under the control of the Thy-1 promoter. Mice were identified by earmarks and numbered accordingly. Animals were randomly assigned to five experimental groups: control, untreated, robot, toxin and combined treatment (toxin and combined treatment mice are a subgroup of those used in [1]). Each group contained comparable numbers of male and female mice (weighing approximately 25g). The age of mice was consistent between the groups (ranging from 3 to 4 month).

*Surgical Procedures*

All surgical procedures were performed under Isoflurane anesthesia (3% induction, 1.5% maintenence, in 1.5L/min oxygen). The animals (apart from the control mice) were placed into a stereotaxic apparatus (Stoelting, Wheat Lane, Wood Dale, IL 60191) and, after removing the skin over the skull and the periosteum, the primary motor cortex (M1) was identified (stereotaxic coordinates 1,75 lateral, 0.5 anterior to bregma). Five minutes after intraperitoneal injection of Rose Bengal (0.2 ml, 10 mg/ml solution in Phosphate Buffer Saline (PBS); Sigma Aldrich, St. Louis, Missouri, USA), white light from an LED lamp (CL 6000 LED, Carl Zeiss Microscopy, Oberkochen, Germany) was focused with a 20X objective (EC Plan Neofluar NA 0.5, Carl Zeiss Microscopy, Oberkochen, Germany) and used to illuminate the M1 for 15 min to induce unilateral stroke in the right hemisphere. We choose a photothrombotic stroke model as a non invasive technique to induce a targeted ischemic stroke highly reproducible. Lesion volume 30 days after photothrombosis was comparable between animals ($1.2 \pm 0.1mm^2$, average ± SEM). Botulinum Neurotoxin E (BoNT/E) injections in toxin and combined treatment mice were performed during the same surgical session of the photothrombotic lesions. We used a dental drill to create a small craniotomy over M1 of the healthy hemisphere (ML: -1.75; RC:+0.5). Then 500 nL of BoNT/E were delivered in two separate injections. A cover glass and an aluminum headpost were attached to the intact skull using transparent dental cement (Super Bond, C&S). Afterwards, the animals were placed in their cages until full recovery.

*Motor Training Protocol on the M-Platform*

Before the first imaging session each mouse was allowed to become accustomed to the apparatus. The animals were trained by means of the M-Platform, which is a robotic system that encourages mice to perform a retraction movement of their left forelimb [2, 1]. The task consisted of up to 15 cycles of passive extension of the affected forelimb followed by its active retraction triggered by an acoustic cue. The time course of one individual training cycle is detailed in Table 1. All groups performed at least four weeks (20 sessions) of daily sessions; in addition, 5 out of 8 robot mice were also recorded for one week before stroke (5 sessions). The M-Platform was designed to allow mice in all conditions (before stroke, right after stroke, and during the weeks under all treatments) to easily perform the motor task from the very first session by applying similar forces. For the same reason, this robotic device is not suitable to evaluate post-stroke functional impairment.

*Corresponding author
[1] These authors contributed equally to this work

| Status | |
|---|---|
| 0 | linear actuator positions forelimb at 10 mm from resting position (passive maximum extension) |
| 1 | forelimb remains in extended position (0.50s) |
| 2 | acoustic tone (1.00V, 0.50s) signals beginning of task |
| 3 | mouse is allowed to perform task of pulling handle back to resting position |
| 4 | different acoustic tone (3.00V, 1.00s) marks end of task |
| 5 | waiting time before reward supply (0.50s) |
| 6 | supply of liquid reward for successful execution of task (0.30s) |
| 7 | waiting time (2.00s) to allow mouse to drink reward before next task |

Table 1: Time course of training cycle. Each line corresponds to a different value of the status variable.

*Wide-Field Fluorescence Microscopy*

The custom-made wide-field imaging setup [3, 4, 5] was equipped with a 505 nm LED (M505L3 Thorlabs, New Jersey, United States) light was deflected by a dichroic filter (DC FF 495-DI02 Semrock, Rochester, New York USA) on the objective (2.5x EC Plan Neofluar, NA 0.085, Carl Zeiss Microscopy, Oberkochen, Germany). Then a 20x objective (LD Plan Neofluar, 20x/0.4 M27, Carl Zeiss Microscopy, Oberkochen, Germany) was used to demagnify the image onto a high-speed complementary metal-oxide semiconductor (CMOS) sensor (OrcaFLASH 4.0, Hamamatsu Photonics, NJ, USA). The fluorescence signal was selected by a band pass filter (525/50 Semrock, Rochester, New York USA) and images (100 x 100 pixels, pixel size 60 $\mu$m) were acquired at 25 Hz. Accordingly, signals from every pixel reflect the activity of hundreds if not thousands of neurons.

*Schallert cylinder test*

For the evaluation of motor performances mice were placed in a Plexiglas cylinder (7,5 cm diameter, 16 cm height) and, after two minutes of acclimatization, recorded for five minutes by a webcam placed below the cylinder. We analyzed mice spontaneous forelimb use through six time points (before stroke and two days after, and once a week during the four week rehabilitative period). Videos were monitored frame by frame and the spontaneous use of both forelimbs was assessed during exploration of the walls, by counting the number of contacts performed by the paws of the animal. For each wall exploration, the last paw that left and the first paw that contacted the wall or the ground were assessed by an experimenter who was blind to the experimental group. In order to quantify forelimb-use asymmetry displayed by the animal, an asymmetry index was computed, according to Lai et al. 2015 [6]:

$$A = \frac{C_{ipsi} - C_{contra}}{C_{ipsi} + C_{contra}} * 100 \quad (1)$$

where $C_{ipsi}$ and $C_{contra}$ refer, respectively, to the number of contacts performed with the limb ipsilateral and contralateral to the lesioned hemisphere.

*Signal processing and data analysis*

*Preprocessing*

Data acquired during each recording session (one mouse, one day, see Fig. 1b) was processed offline using custom routines implemented in Python (Python Software Foundation) and Matlab (MathWorks). Each such dataset consisted of up to 15 cycles of active retraction movements on a slide triggered by passively actuated contralesional forelimb extensions. To ensure the consistency of the field of view across sessions and across mice, each frame of the fluorescence data was offline registered by aligning each frame to two reference points (corresponding to bregma and lambda) that were previously marked on the glass window during the surgery procedure. For the 2D fluorescence data, masking the region of interest and spatial downsampling by a factor 3 for both rows and columns resulted in calcium activity matrices of 12 x 21 pixels. Spatial average over all pixels yielded the mean calcium activity. In parallel, the force applied to the slide by the mouse and the discrete status of the slide were recorded. Using samplings with a time step of 40 ms and acquisition times of up to 400 seconds this yielded recordings with at most 10000 data points. The calcium traces were detrended via subtraction of a moving average of order 75 (three seconds) and, in order to yield a better time resolution, upsampled by a factor 20.

*Event detection*

Next, within all of these traces we identified the times of the most relevant discrete events. For the status (Fig. 2a) we marked the transition from level 3 to level 4 which corresponds to the completion of the forelimb retraction by the active movement of the mouse upon which the animal received its reward (reward pulling event). For the force (Fig. 2b), the mean calcium (Fig. 2c) and the individual calcium traces of all the pixels the events are the high-amplitude peaks that can easily be recognized. As event times we used the upwards crossings of a threshold T which in each of these cases was defined in a data-adaptive manner according to $T = mean(x) + t * std(x)$. The free parameter $t$ was set to 1.5 for the force and 1.7 for all the calcium traces. In the slower calcium traces, in order to avoid a double detection due to noise, we discarded all events that succeeded the previous event by less than a minimum inter-event interval of 25 data points (one second).

*SPIKE-Order*

The events (from now on called spikes) of all the pixels can be represented best in a rasterplot like the one shown in Fig. 2d. The next important step was to identify the global events that correspond to the events of the mean calcium trace. To this aim, we used the cSPIKE-implementation [7] of the SPIKE-Order approach recently proposed in [8] (for detailed mathematical definitions of all the underlying quantities please refer to the Appendix "SPIKE-Order method". The original proposal was designed for rather clean data with well-defined global events. These conditions hold for most of our datasets as well, however, we added a few tailor-made denoising steps that addressed the rare instances of increased noisiness that we observed in some of the datasets.

The procedure consisted of six steps: in an initial denoising step, we filtered out all spikes of individual pixels that were not within 1 second of a mean calcium event and thus were certainly not part of global events. Secondly, we applied the coincidence detection first introduced for the bivariate measure *event synchronization* [9]. This criterion paired spikes in such a way that every spike was matched with at most one spike in each of the other pixels. Here we combined the original adaptive approach with a maximum allowed distance between spikes of 2.5 seconds. Next, we used the symmetric and multivariate measure SPIKE-Synchronization C [10] to quantify for each spike the fraction of other pixels for whom a matching spike could be found. By setting a threshold value $C_{thr} = 0.75$ we only took into account spikes which were coincident with spikes in at least three quarters of the other pixels, all other spikes were filtered out as background noise.

In a fourth step, we applied the SPIKE-Order $D$ [8] which evaluates the temporal order of the spikes by quantifying for each spike the net-fraction of spikes of other pixels this spike is leading (positive value) or following (negative value). Based on the time profile we identified start and end spikes of global events by tracking the jumps from a negative local minimum (last spike of previous event) to a positive local maximum (first spike of current event). In one further denoising step we discarded split events and eliminated outlier spikes by using a maximum distance between consecutive spikes of 0.15 seconds and thereby kept only continuous global events. The final step used in the visualization of the spike trains in Fig. 2d involved the Synfire Indicator [8], a scalar measure which quantifies to what degree the spatiotemporal propagation patterns of the global events are consistent with each other. Optimization of this indicator was used to sort the spike trains / pixels from overall leader to overall follower. Here, overall means that we take into account all global events at the same time. The result is that the first spike trains contain mostly leading spikes, whereas the last spike trains consist largely of trailing spikes.

*Categorization of events*

Next, we divided the global events into several types using the following three-level categorization scheme (the corresponding branching structure is shown in Section Methods): First, we separated all the global events that are not associated with a force event (non-Force, **nF**). For this we demanded that there is no force event in the interval [1 second before, 0.75 seconds after] the matching calcium event. The window was slightly asymmetric to account for the fact that typically Force were observed a bit earlier than mean calcium events. The remaining force events (**F**) were further subdivided into events that occur during the passive extension of the arm by the slide (Passive, **Pass**) and events that occur outside that window (Active, **Act**). In the passive events the mouse applied force to resist the forelimb extension movement of the robot, whereas the active events were the ones where the force was applied during an active retraction movement (when the status variable was set to 3, i.e. between the Go cue and the completion of the task). Finally, among the active events we distinguished between events which were not completed and thus not rewarded (non-Reward Pulling, **nRP**) and events which lead to a completion of the forelimb retraction and therefore were rewarded with milk (Reward Pulling, **RP**). The categorization criterion was the occurrence of a transition from status 3 to status 4 within [0.75 seconds before, 0.75 seconds after] a calcium event. This window was symmetric, since the observed temporal distribution of status events was symmetric with respect to the mean calcium events.

*Three propagation indicators: Duration, Angle, Smoothness*

For all global events, the event time was defined as the average time of all the spikes within the event and our first propagation indicator, the event duration, was defined as time from the first to the last spike of the event. To calculate the other two propagation indicators, angle and smoothness, we first generated the propagation matrix by mapping the color-coded relative order of the spikes onto the pixels of the 2D-recording plane (compare Fig. 2f). Next, we applied singular value decomposition (SVD, [11]) which searches for spatial patterns by decomposing the propagation matrix $P$ into three simple transformations: a rotation $U$, a scaling $\Sigma$ along the rotated coordinate axes and a second rotation $V^T$.

The rotations $U$ and $V^T$ are orthonormal matrices and $\Sigma$ is a diagonal matrix containing in its diagonal the singular values $\sigma_i$ of $P$. By backprojecting the sorted singular values one at a time

$$\Sigma_1 = \begin{bmatrix} \sigma_1 & & \\ & 0 & \\ & & 0 \\ & & & \ddots \end{bmatrix} \qquad \Sigma_2 = \begin{bmatrix} 0 & & \\ & \sigma_2 & \\ & & 0 \\ & & & \ddots \end{bmatrix}$$

we could obtain various projections of the original propagation matrix

$$P_1 = U\Sigma_1 V^T \qquad\qquad P_2 = U\Sigma_2 V^T.$$

The mean gradients with respect to column ($c$) and row ($r$) increments of the first two projections were calculated as

$$\begin{cases} g_1^c = \mathbb{E}(-\frac{\partial P_1}{\partial c}) \\ g_1^r = \mathbb{E}(-\frac{\partial P_1}{\partial r}) \end{cases} \qquad\qquad \begin{cases} g_2^c = \mathbb{E}(-\frac{\partial P_2}{\partial c}) \\ g_2^r = \mathbb{E}(-\frac{\partial P_2}{\partial r}) \end{cases}$$

with $\mathbb{E}$ denoting the average across pixels while the sign (-) is defined by the directionality in the matrix $P$ going from leader (+1) to follower (-1). The main propagation directions, along the column and row directions,

$$\begin{cases} v^c = \sigma_1 g_1^c + \sigma_2 g_2^c \\ v^r = \sigma_1 g_1^r + \sigma_2 g_2^r \end{cases}$$

were calculated from the weighted average of the mean gradients of the first two projections, with the singular values as weights. Our second propagation indicator, the angle

$$\alpha = \arctan\left(\frac{v^c}{v^r}\right)$$

was defined relative to the horizontal axis.

Finally, our third propagation indicator, the smoothness $S$, quantified how well the second order approximation, the weighted sum of the projections of only the first two singular values, captures the full spatiotemporal pattern obtained by considering all singular values $\sigma_i$. Smoothness is defined as the relative weight of the first two approximations

$$S = \frac{\sigma_1^2 + \sigma_2^2}{\sum_i \sigma_i^2}. \tag{2}$$

*Statistical Tests*

Considering the daily averages of the events as the unit observations we performed a separate analysis for each propagation indicator, event type and condition: healthy (Fig. 4), acute phase (Fig. 5), comparison of treatments (Fig. 6), pre-stroke (Fig. 2) and untreated acute phase (Fig. 3). For smoothness, duration and the asymmetry index (Fig. 4) we considered intra-subject temporal correlation by means of mixed-effect models, while for the propagation angle we adopted the Von-Mises distribution to model the circular characteristic of this indicator.

Using the *R* library *lme4* [12] for inference, we initialized the mixed models including all possible fixed effects, their interactions, a random intercept per subject, a random slope for the daily trend and the random intercept-random slope covariance. Afterwards, we carried out a backward selection with consecutive likelihood-ratio tests. Here we used the function *step* of the *lmerTest* library [13] in order to select a parsimonious feasible model without removing relevant effects that could decrease the *type I error* rate and increase the statistical power [14]. Once the model parameters were selected, we tested the differences between the least-squares means with the *difflsmeans* function of the *lmerTest* library and then corrected the results with the Holm-Bonferroni correction.

For every test we assessed both the normality of residuals and the homogeneity of variance assumptions before reporting the results. We used the Kolmogorov-Smirnov test to check the residual distribution and the Breusch-Pagan test to check the homoscedasticity. If normality assumption did not hold, we adopted a Box-Cox transform of the dependent variable [15]. In case the Breusch-Pagan test revealed a departure from homogeneity of variance, we identified the comparisons that had a significant difference in variance and reported the results, again correcting the p-values with the Holm-Bonferroni method.

For the propagation angle, we tested the differences in circular variance with multiple Bartlett tests [*equal.kappa.test* function in the *circular* library, 16]. Again, we adopted the Holm-Bonferroni correction to account for multiple comparison bias and we assessed the assumption of Von-Mises distribution with the Watson test [17]. We only reported results of those comparisons for which the Von-Mises distribution held.

### Appendix: SPIKE-Order method

The central method of our study is the SPIKE-Order approach which we use to identify global events and to track the propagation patterns within these events by sorting the spikes from leader to follower. Here we present the more detailed mathematical definitions:

*Adaptive Coincidence Detection*

Analyzing leader-follower relationships in a spike train set requires a criterion that determines which spikes should be compared against each other. Here we use the adaptive coincidence criterion first proposed in [9]. This coincidence detection is scale- and parameter-free since the maximum time lag $\tau_{ij}^{(m,n)}$ up to which two spikes $t_i^{(m)}$ and $t_j^{(n)}$ of spike trains $m, n = 1, ..., N$ (with $N$ denoting the number of spike trains) are considered to be synchronous is adapted to the local firing rates according to

$$\begin{aligned} \tau_{ij}^{(m,n)} = \min\{&t_{i+1}^{(m)} - t_i^{(m)}, t_i^{(m)} - t_{i-1}^{(m)}, \\ &t_{j+1}^{(n)} - t_j^{(n)}, t_j^{(n)} - t_{j-1}^{(n)}\}/2. \end{aligned} \tag{3}$$

*SPIKE-Synchronization*

Following [10], we apply the adaptive coincidence criterion in a multivariate context by defining for each spike $i$ of any spike train $n$ and for each other spike train $m$ a coincidence indicator

$$C_i^{(n,m)} = \begin{cases} 1 & \text{if } \min_j(|t_i^{(n)} - t_j^{(m)}|) < \tau_{ij}^{(n,m)} \\ 0 & \text{otherwise.} \end{cases} \tag{4}$$

which is either one or zero depending on whether this spike is part of a coincidence with a spike of spike train $m$ or not. This results in an unambiguous spike matching since any spike can at most be coincident with one spike (the nearest one) in the other spike train.

Subsequently, for each spike of every spike train a normalized coincidence counter

$$C_i^{(n)} = \frac{1}{N-1} \sum_{m \neq n} C_i^{(n,m)} \quad (5)$$

is obtained by averaging over all $N-1$ bivariate coincidence indicators involving the spike train $n$.

In order to obtain a single multivariate SPIKE-Synchronization profile we pool the coincidence counters of all the spikes of every spike train:

$$\{C(t_k)\} = \bigcup_n \{C_{i(k)}^{(n(k))}\}, \quad (6)$$

where we map the spike train indices $n$ and the spike indices $i$ into a global spike index $k$ denoted by the mapping $i(k)$ and $n(k)$.

With $M$ denoting the total number of spikes in the pooled spike train, the average of this profile

$$S_C = \begin{cases} \frac{1}{M}\sum_{k=1}^{M} C(t_k) & \text{if } M > 0 \\ 1 & \text{otherwise} \end{cases} \quad (7)$$

yields SPIKE-Synchronization, the overall fraction of coincidences. It reaches one if and only if each spike in every spike train has one matching spike in all the other spike trains (or if there are no spikes at all), and it attains the value zero if and only if the spike trains do not contain any coincidences.

*SPIKE-Order*

While SPIKE-Synchronization is invariant to which of the two spikes within a coincidence is leading and which is following, the temporal order of the spikes is taken into account by the two indicators SPIKE-Order and Spike Train Order.

The bivariate anti-symmetric SPIKE-Order indicators

$$\begin{aligned} D_i^{(n,m)} &= C_i^{(n,m)} \cdot \text{sign}(t_{j'}^{(m)} - t_i^{(n)}) \\ D_{j'}^{(m,n)} &= C_{j'}^{(m,n)} \cdot \text{sign}(t_i^{(n)} - t_{j'}^{(m)}) = -D_i^{(n,m)} \end{aligned} \quad (8)$$

where the index $j'$ is defined from the minimum in Eq. 4 as $j' = \arg\min_j(|t_i^{(1)} - t_j^{(2)}|)$, assign to each spike either a 1 or a $-1$ depending on whether the respective spike is leading or following a coincident spike in the other spike train.

SPIKE-Order distinguishes leading and following spikes, and is thus used for color-coding the individual spikes on the leader to follower scale. But it can also be employed to sort the *spike trains* based on a pairwise analysis. For this we use the cumulative SPIKE-Order matrix

$$D^{(n,m)} = \sum_i D_i^{(n,m)}. \quad (9)$$

This anti-symmetric matrix sums up the orders of coincidences from the respective pair of spike trains only and quantifies how much spike train $n$ is leading spike train $m$. Hence if $D^{(n,m)} > 0$ spike train $n$ is leading $m$, while $D^{(n,m)} < 0$ means $m$ is leading $n$. If the current spike train order is consistent with the synfire property (i.e., it displays consistent repetitions of the same global propagation pattern), we thus expect that $D^{(n,m)} > 0$ for $n < m$ and $D^{(n,m)} < 0$ for $n > m$. Therefore, we construct the overall SPIKE-Order as

$$D_< = \sum_{n<m} D^{(n,m)}, \quad (10)$$

i.e. the sum over the upper right tridiagonal part of the matrix $D^{(n,m)}$.

*Synfire Indicator*

After normalizing by the overall number of possible coincidences, we arrive at the definition of the Synfire Indicator:

$$F = \frac{2D_<}{(N-1)M}. \quad (11)$$

This measure quantifies to what degree coinciding spike pairs with correct order prevail over coinciding spike pairs with incorrect order, or in other words, to what extent the spike trains in their current order resemble a synfire pattern. Conversely, the maximization of the Synfire Indicator as a function of the spike train order within a set of spike trains can be used to sort spike trains from leader to follower such that the set comes as close as possible to a synfire pattern. Denoting the Synfire Indicator for any given spike train index permutation $\varphi(n)$ as $F_\varphi$, the optimal (**s**orted) order $\varphi_s$ is the one resulting in the maximal overall Synfire Indicator $F_s = F_{\varphi_s}$:

$$\varphi_s : F_{\varphi_s} = \max_\varphi\{F_\varphi\} = F_s. \quad (12)$$

Whereas the Synfire Indicator $F_\varphi$ for any spike train order $\varphi$ is normalized between $-1$ and 1, the optimized Synfire Indicator $F_s$ can only attain values between 0 and 1. A perfect synfire pattern results in $F_s = 1$, while sufficiently long Poisson spike trains without any synfire structure yield $F_s \approx 0$. For details on the optimization procedure, please refer to [8].

### References

### References

[1] A. L. Allegra Mascaro, E. Conti, S. Lai, A. P. Di Giovanna, C. Spalletti, C. Alia, A. Panarese, A. Scaglione, L. Sacconi,

| Panel | Indicator | Event type | Group | Diff. type | p-value | |
|---|---|---|---|---|---|---|
| c | Duration | Act-Pass | Control | Variance | 0.002 | $\star\star$ |
| e | Smoothness | F-nF | | | 0.001 | $\star\star$ |
| | | RP-nRP | | | 0.001 | $\star\star$ |
| g | Angle | F-nF | | | $10^{-10}$ | $\star\star\star$ |
| | | Act-Pass | | | $10^{-9}$ | $\star\star\star$ |
| | | RP-nRP | | | $10^{-8}$ | $\star\star\star$ |

Table 2: P-values for Fig. 4

| Panel | Indicator | Event type | Group | Diff. type | p-value | |
|---|---|---|---|---|---|---|
| a | Duration | | Control - Acute stroke | Mean | 0.003 | ** |
| b | | F | Control - Acute stroke | | 0.026 | * |
| | | Act-Pass | Control | | 0.042 | * |
| | | Act | Control - Acute stroke | | 0.034 | * |
| | | Pass | Control - Acute stroke | | 0.035 | * |
| | | RP | Control - Acute stroke | | 0.014 | * |
| | | nRP | Control - Acute stroke | | 0.028 | * |
| c | Smoothness | | Control - Acute stroke | | 0.007 | ** |
| d | | F - nF | Control | | $10^{-10}$ | *** |
| | | F | Control - Acute stroke | | 0.004 | ** |
| | | Act | Control - Acute stroke | | 0.01 | * |
| | | Pass | Control - Acute stroke | | 0.002 | ** |
| | | RP-nRP | Control | | 0.008 | ** |
| | | RP | Control - Acute stroke | | 0.005 | ** |
| f | Angle | F - nF | Control | Variance | $10^{-7}$ | $\star\star\star$ |
| | | | Acute stroke | | $10^{-5}$ | $\star\star\star$ |
| | | Act - Pass | Control | | $10^{-4}$ | $\star\star\star$ |
| | | | Acute stroke | | 0.046 | $\star$ |

Table 3: P-values for Fig. 5


S. Micera, M. Caleo, F. S. Pavone, Combined rehabilitation promotes the recovery of structural and functional features of healthy neuronal networks after stroke, Cell Rep. 28 (13) (2019) 3474–3485.e6.

[16] C. Agostinelli, U. Lund, R package `circular`: Circular Statistics (version 0.4-93), CA: Department of Environmental Sciences, Informatics and Statistics, Ca' Foscari University, Venice, Italy.

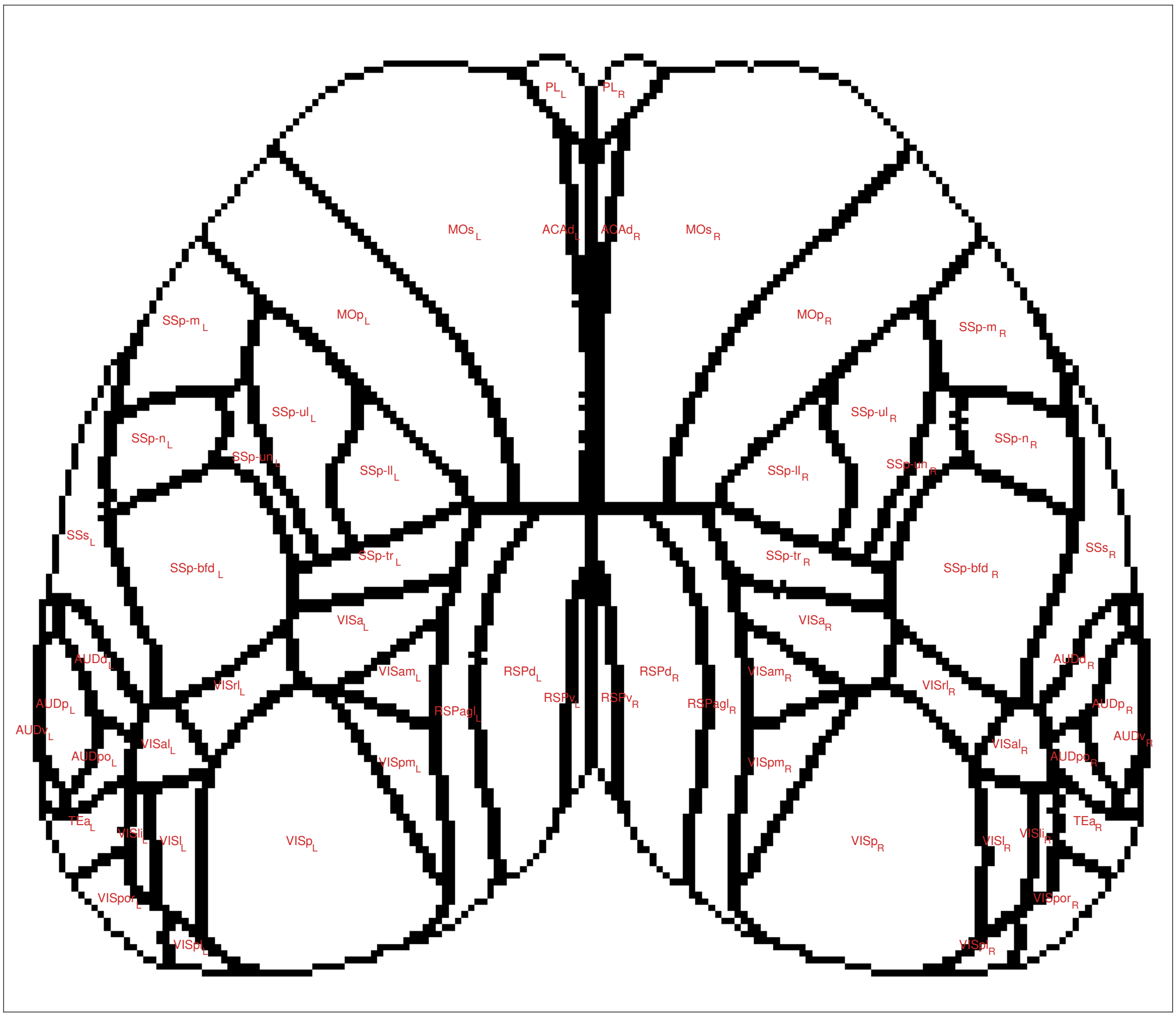


| Acronym | Name | Acronym | Name |
|---|---|---|---|
| MOp | Primary motor area | VISam | Anteromedial visual area |
| MOs | Secondary motor area | VISl | Lateral visual area |
| SSp-n | Primary somatosensory area | VISp | Primary visual area |
| SSp-bfd | Primary somatosensory area | VISpl | Posterolateral visual area |
| SSp-ll | Primary somatosensory area | VISpm | posteromedial visual area |
| SSp-m | Primary somatosensory area | VISli | Laterointermediate area |
| SSp-ul | Primary somatosensory area | VISpor | Postrhinal area |
| SSp-tr | Primary somatosensory area | ACAd | Anterior cingulate area |
| SSp-un | Primary somatosensory area | PL | Prelimbic area |
| SSs | Supplemental somatosensory area | RSPagl | Retrosplenial area |
| AUDd | Dorsal auditory area | RSPd | Retrosplenial area |
| AUDp | Primary auditory area | RSPv | Retrosplenial area |
| AUDpo | Posterior auditory area | VISa | Anterior area |
| AUDv | Ventral auditory area | VISrl | Rostrolateral visual area |
| VISal | Anterolateral visual area | TEa | Temporal association areas |

Figure 1: Standard atlas of brain regions and their acronyms.

UL: Department of Statistics, California Polytechnic State University, San Luis Obispo, California, USA (2017).
URL `https://r-forge.r-project.org/projects/circular/`

[17] M. A. Stephens, Use of the kolmogorov–smirnov, cramer–von mises and related statistics without extensive tables, Journal of the Royal Statistical Society: Series B (Methodological) 32 (1) (1970) 115–122.

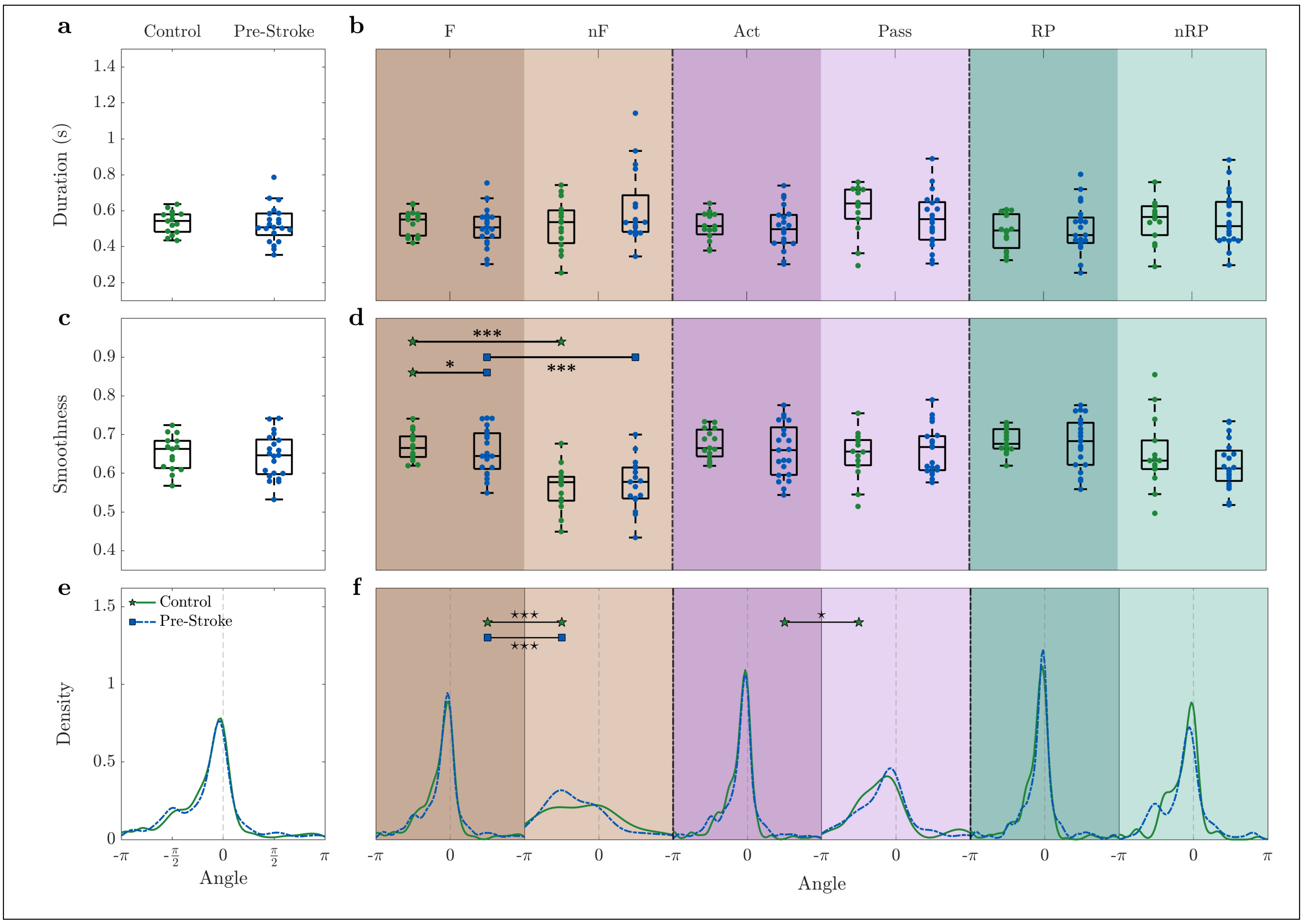


Figure 2: As expected, the pre-stroke condition shows the same behavior as the control group for all three propagation indicators: **(a-b)** duration, **(c-d)** smoothness, and **(e-f)** angle. — Duration and angle are weighted by smoothness. Markers in (a-d) refer to the average value per day. Within each box in (a-d), the central mark indicates the median, and the bottom and top edges of the box indicate the $25^{th}$ and $75^{th}$ percentiles, respectively. Control group n=3 mice, pre-stroke group n=5 mice.

| Panel | Indicator | Event type | Group | Diff. type | p-value | |
|---|---|---|---|---|---|---|
| a | Duration | | Untreated - Combined | Mean | $10^{-4}$ | *** |
| | | | Robot - Combined | | $10^{-5}$ | *** |
| | | | Toxin - Combined | | 0.006 | ** |
| b | | F | Untreated - Combined | | 0.001 | ** |
| | | | Robot - Combined | | $10^{-5}$ | *** |
| | | | Toxin - Combined | | 0.035 | * |
| | | nF | Untreated - Combined | | $10^{-6}$ | *** |
| | | | Robot - Combined | | $10^{-7}$ | *** |
| | | | Toxin - Combined | | 0.002 | ** |
| | | F-nF | Untreated | | 0.037 | * |
| | | | Robot | | 0.020 | * |
| | | Act | Untreated - Combined | | 0.002 | ** |
| | | | Robot - Combined | | $10^{-5}$ | *** |
| | | Pass | Untreated - Combined | | $10^{-4}$ | *** |
| | | | Robot - Combined | | $10^{-4}$ | *** |
| | | | Toxin - Combined | | 0.01 | * |
| | | Act-Pass | Toxin | | 0.015 | * |
| | | RP | Untreated - Combined | | 0.006 | ** |
| | | | Robot - Combined | | $10^{-5}$ | *** |
| | | nRP | Untreated - Combined | | $10^{-4}$ | *** |
| | | | Robot - Combined | | $10^{-5}$ | *** |
| | | | Toxin - Combined | | 0.003 | ** |
| | | RP-nRP | Toxin | | 0.006 | ** |
| c | Smoothness | | Untreated - Toxin | | 0.029 | * |
| | | | Untreated - Combined | | $10^{-5}$ | *** |
| | | | Robot - Combined | | $10^{-4}$ | *** |
| | | | Toxin - Combined | | 0.003 | ** |
| d | | F | Untreated - Combined | | $10^{-5}$ | *** |
| | | | Robot - Combined | | $10^{-4}$ | *** |
| | | | Toxin - Combined | | 0.019 | * |
| | | nF | Untreated - Combined | | $10^{-5}$ | *** |
| | | | Robot - Combined | | $10^{-4}$ | *** |
| | | | Toxin - Combined | | $10^{-4}$ | *** |
| | | F-nF | Robot | Variance | 0.007 | ⋆⋆ |
| | | | Toxin | Mean | 0.015 | * |
| | | Act | Untreated - Combined | | $10^{-4}$ | *** |
| | | | Robot - Combined | | $10^{-4}$ | *** |
| | | | Toxin - Combined | | 0.023 | * |
| | | Pass | Untreated - Combined | | $10^{-5}$ | *** |
| | | | Robot - Combined | | $10^{-4}$ | *** |
| | | | Toxin - Combined | | 0.013 | * |
| | | RP | Untreated - Combined | | $10^{-5}$ | *** |
| | | | Robot - Combined | | $10^{-5}$ | *** |
| | | | Toxin - Combined | | 0.005 | ** |
| | | nRP | Untreated - Combined | | $10^{-5}$ | *** |
| | | | Robot - Combined | | $10^{-4}$ | *** |
| | | | Toxin - Combined | | 0.010 | * |
| e | Angle | | Untreated - Robot | Variance | 0.018 | ⋆ |
| | | | Robot - Toxin | | 0.005 | ⋆⋆ |
| f | | F | Untreated - Robot | | 0.037 | ⋆ |
| | | | Robot - Toxin | | 0.003 | ⋆⋆ |
| | | F-nF | Robot | | $10^{-10}$ | ⋆⋆⋆ |
| | | Act | Untreated - Robot | | 0.001 | ⋆⋆ |
| | | | Robot - Toxin | | $10^{-4}$ | ⋆⋆⋆ |
| | | RP | Untreated - Robot | | $10^{-5}$ | ⋆⋆⋆ |
| | | | Robot - Toxin | | $10^{-4}$ | ⋆⋆⋆ |



Table 4: P-values for Fig. 6

| Panel | Indicator | Event type | Group | Diff. type | p-value | |
|---|---|---|---|---|---|---|
| d | Smoothness | F - nF | Control | Mean | $10^{-5}$ | *** |
| | | | Pre-stroke | | $10^{-4}$ | *** |
| | | F | Control - Pre-stroke | | 0.049 | * |
| f | Angle | F - nF | Control | Variance | $10^{-4}$ | $\star\star\star$ |
| | | | Pre-stroke | | $10^{-9}$ | $\star\star\star$ |
| | | Act - Pass | Control | | 0.012 | $\star$ |

Table 5: P-values for Fig. 2

| Panel | Indicator | Event type | Group | Diff. type | p-value | |
|---|---|---|---|---|---|---|
| c | Smoothness | | Acute stroke - Untreated | Mean | 0.005 | ** |
| d | Smoothness | Act - Pass | Acute stroke | | 0.038 | * |
| | | Act | Acute stroke - Untreated | | 0.023 | * |
| | | Pass | | | 0.016 | * |
| f | Angle | F - nF | Acute stroke | Variance | $10^{-5}$ | $\star\star\star$ |
| | | | Untreated | | 0.042 | $\star$ |

Table 6: P-values for Fig. 3

| Indicator | Group | Diff. type | p-value | |
|---|---|---|---|---|
| Asymmetry index | Untreated - Combined | Mean | $10^{-4}$ | *** |
| | Robot - Combined | | $10^{-5}$ | *** |
| | Toxin - Combined | | $10^{-6}$ | *** |

Table 7: P-values for Fig. 4

| Panel | Indicator | Event type | Group | Diff. type | p-value | |
|---|---|---|---|---|---|---|
| a | Duration | | Control - Combined | Mean | $10^{-4}$ | *** |
| | | | Robot - Combined | | $10^{-6}$ | *** |
| b | | F | Control - Combined | | 0.002 | ** |
| | | | Robot - Combined | | $10^{-5}$ | *** |
| | | nF | Control - Combined | | 0.002 | ** |
| | | | Robot - Combined | | $10^{-5}$ | *** |
| | | Act | Control - Combined | | 0.002 | ** |
| | | | Robot - Combined | | $10^{-4}$ | *** |
| | | Pass | Control - Combined | | $10^{-4}$ | *** |
| | | | Robot - Combined | | $10^{-5}$ | *** |
| | | RP | Control - Combined | | 0.003 | ** |
| | | | Robot - Combined | | $10^{-6}$ | *** |
| | | nRP | Control - Combined | | 0.002 | ** |
| | | | Robot - Combined | | $10^{-6}$ | *** |
| c | Smoothness | | Control - Combined | | 0.025 | * |
| | | | Robot - Combined | | $10^{-4}$ | *** |
| d | | F | Robot - Combined | | 0.001 | ** |
| | | nF | Control - Combined | | 0.018 | * |
| | | | Robot - Combined | | 0.002 | ** |
| | | Act | Control - Robot | | 0.046 | * |
| | | | Robot - Combined | | $10^{-6}$ | *** |
| | | Pass | Control - Combined | | 0.030 | * |
| | | | Robot - Combined | | $10^{-6}$ | *** |
| | | RP | Control - Combined | | 0.029 | * |
| | | | Robot - Combined | | $10^{-5}$ | *** |
| | | nRP | Control - Combined | | 0.014 | * |
| | | | Robot - Combined | | $10^{-4}$ | *** |
| f | Angle | F | Control - Combined | Variance | $10^{-8}$ | $\star\star\star$ |
| | | | Robot - Combined | | $10^{-8}$ | $\star\star\star$ |
| | | Act | Control - Combined | | $10^{-9}$ | $\star\star\star$ |
| | | | Robot - Combined | | $10^{-9}$ | $\star\star\star$ |
| | | Pass | Control - Combined | | 0.027 | $\star$ |
| | | | Robot - Combined | | 0.008 | $\star\star$ |
| | | RP | Control - Combined | | $10^{-9}$ | $\star\star\star$ |
| | | | Robot - Combined | | $10^{-9}$ | $\star\star\star$ |
| | | nRP | Control - Robot | | 0.024 | $\star$ |
| | | | Robot - Combined | | $10^{-5}$ | $\star\star\star$ |

Table 8: P-values for Fig. 5

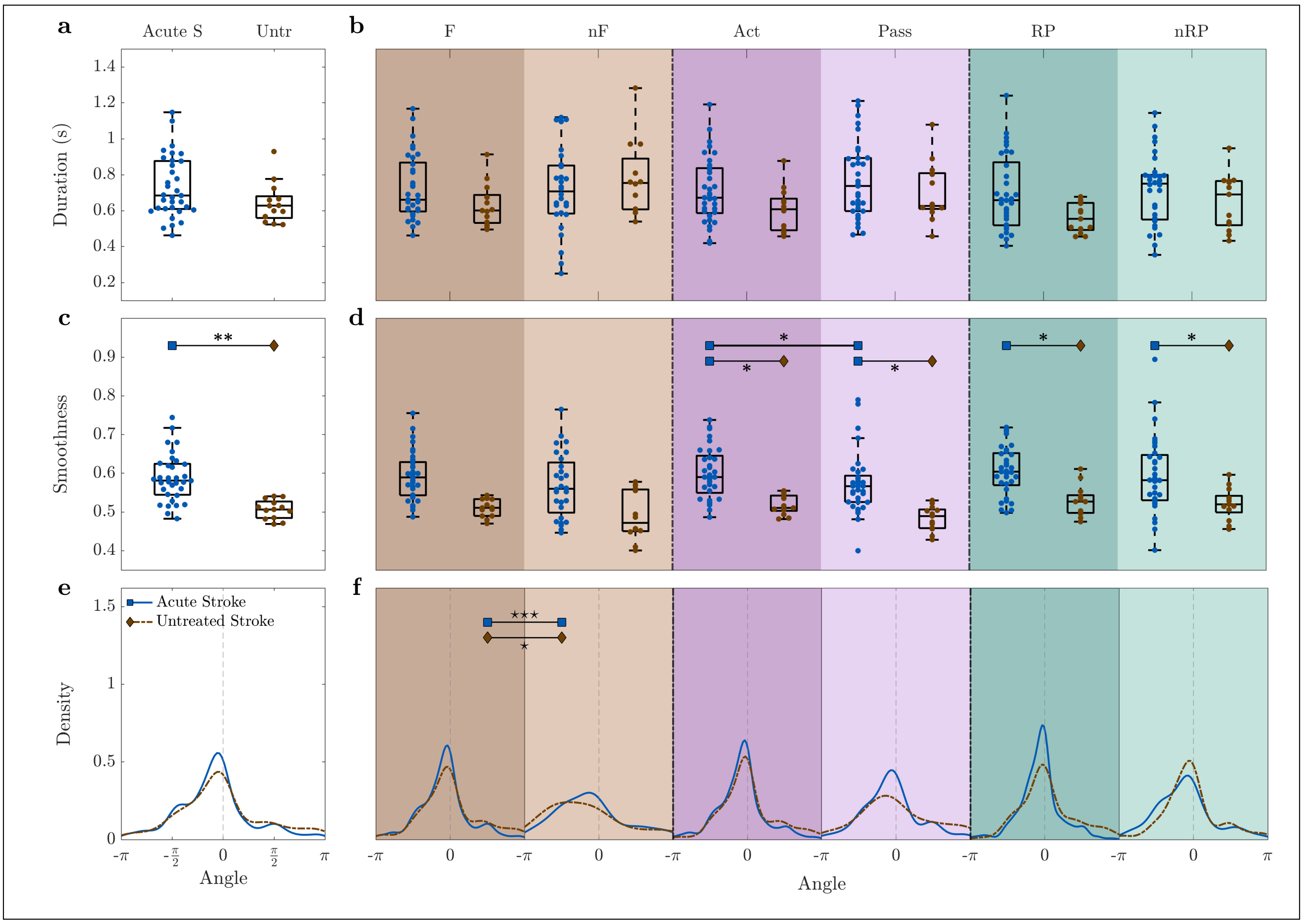


Figure 3: Compared to the acute phase, spontaneously recovered mice (untreated stroke group) present only minor modulations in the propagation indicators: **(a-b)** duration, **(c-d)** smoothness, and **(e-f)** angle. — Duration and angle are weighted by smoothness. Markers in (a-d) refer to the average value per day. Within each box in (a-d), the central mark indicates the median, and the bottom and top edges of the box indicate the $25^{th}$ and $75^{th}$ percentiles, respectively. Control group n=3 mice, pre-stroke group n=5 mice.

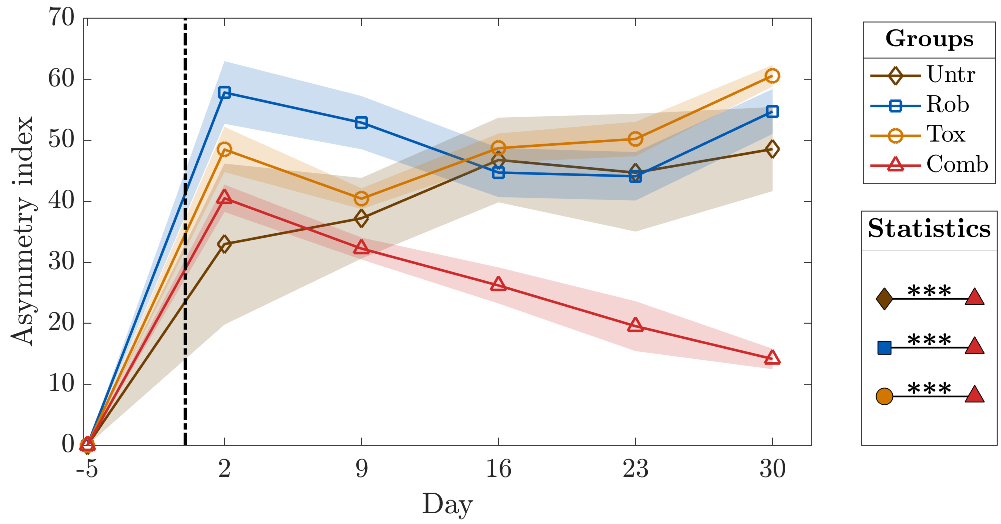


Figure 4: Combined treatment leads to generalized recovery. Asymmetry in the spontaneous forelimb use measured by the Schallert cylinder test once per week and rescaled by the pre-stroke (day -5) value for each mouse. Significant differences are calculated for the last day of recording (day 30). P-values of statistical tests in Table 7.

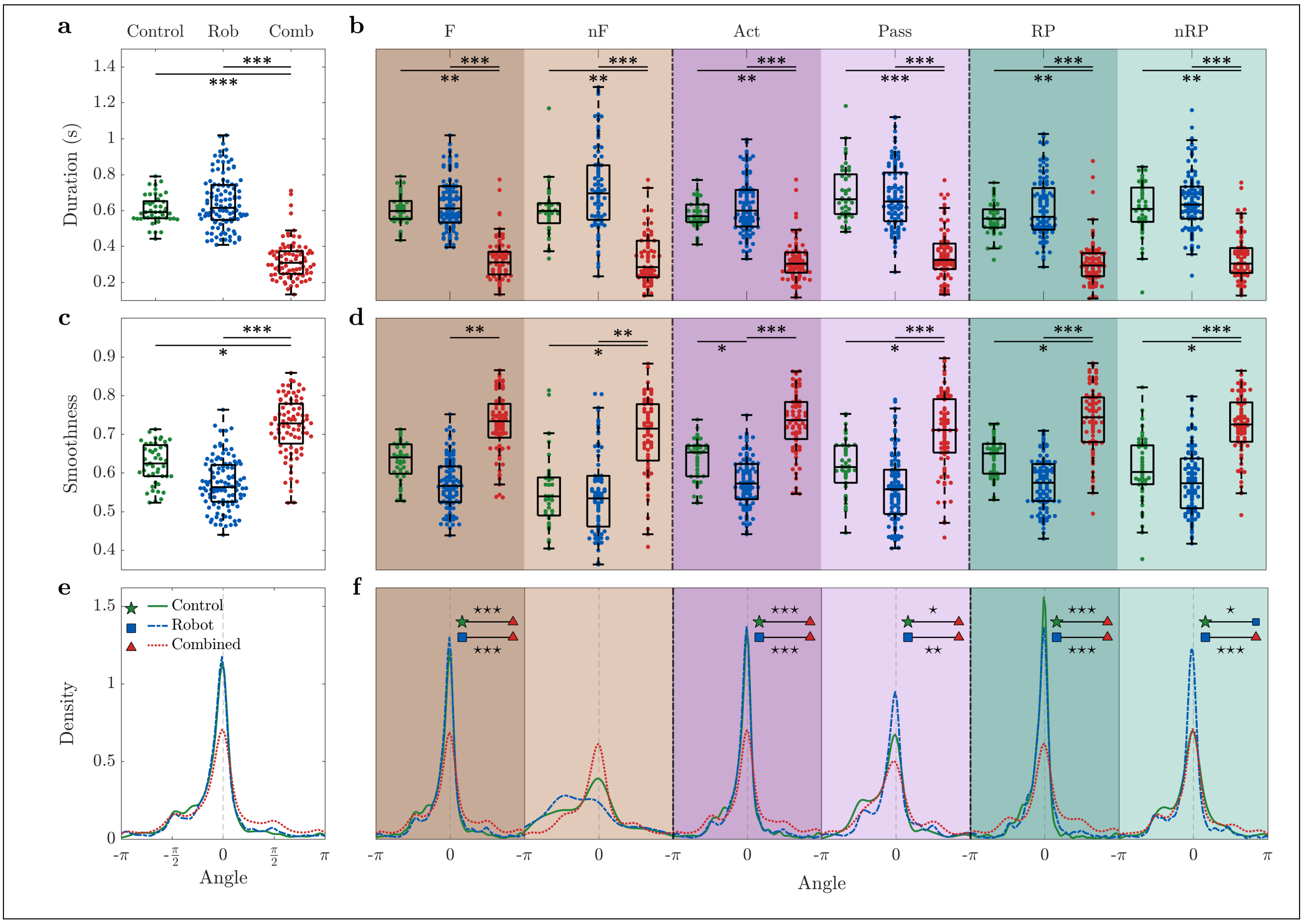


Figure 5: Combined treatment group is characterized by shorter duration and higher smoothness. **(a-b)** For all types of event, combined treatment group events are the shortest. **(c-d)** For all types of event, the smoothness of the combined treatment group is higher than for the robot group. **(e-f)** For the combined treatment group the distribution of the angles does not vary depending on the type of event. — Longitudinal data starting from the second week of recording up to one month after stroke. Duration and angle are weighted by smoothness. Markers in (a-d) refer to the average value per day. Within each box in (a-d), the central mark indicates the median, and the bottom and top edges of the box indicate the $25^{th}$ and $75^{th}$ percentiles, respectively. Control group n=3 mice, robot group n=8 mice, combined treatment group n=6 mice. P-values of statistical tests in Table 8, "⋆" refers to difference in variance and "*" refers to difference in mean.

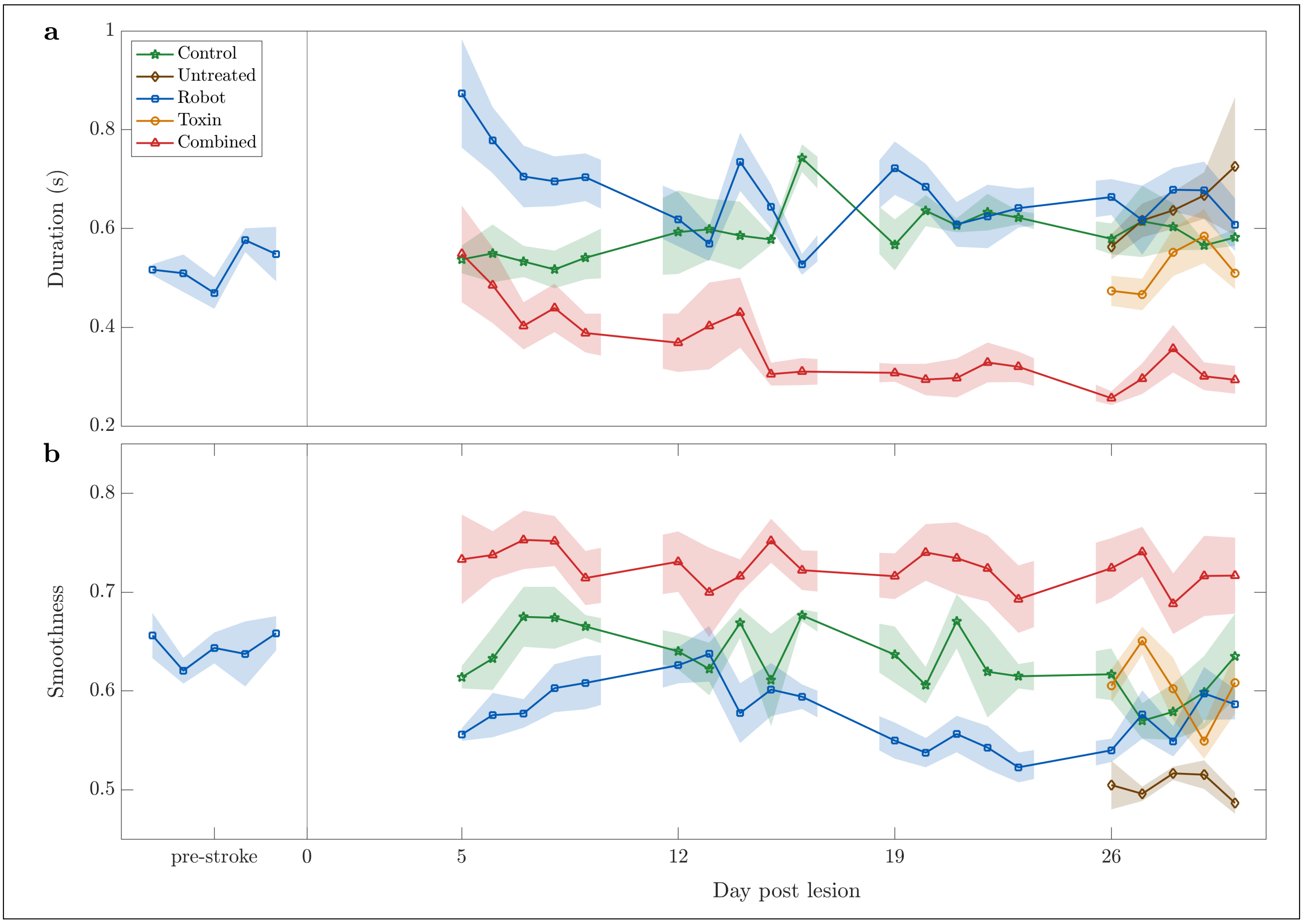


Figure 6: Temporal evolution of duration **(a)** and smoothness **(b)** for all groups. In both cases, the difference between control and acute stroke groups is considerably larger for the first three days. Moreover, the difference in the event duration tends to diminish over days. Event duration is always shorter for the combined treatment group, smoothness is always higher. The control group shows a consistent trend, suggesting that motor training alone has no effect on either duration or smoothness, as already described in the previous sections for healthy mice. After the stroke, the robot group shows a significant variation: the duration is longer and the smoothness is lower; in both cases this variation decreases over time. While the duration reaches again values comparable to healthy mice already after the second week of training, the difference in smoothness seems to oscillate without stabilizing.
